# Granger Causality Inference in EEG Source Connectivity Analysis: A State-Space Approach

**DOI:** 10.1101/2020.10.07.329276

**Authors:** Parinthorn Manomaisaowapak, Anawat Nartkulpat, Jitkomut Songsiri

**Affiliations:** Department of Electrical Engineering, Faculty of Engineering Chulalongkorn University, Bangkok, Thailand 10330

**Keywords:** brain connectivity, state-space models, Granger causality, EEG, group sparse structure

## Abstract

This paper considers a problem of estimating brain effective connectivity from EEG signals using a Granger causality (GC) concept characterized on state-space models. We propose a state-space model for explaining coupled dynamics of the source and EEG signals where EEG is a linear combination of sources according to the characteristics of volume conduction. Our formulation has a sparsity prior on the source output matrix that can further classify active and inactive sources. The scheme is comprised of two main steps: model estimation and model inference to estimate brain connectivity. The model estimation consists of performing a subspace identification and the active source selection based on a group-norm regularized least-squares. The model inference relies on the concept of state-space GC that requires solving a discrete-time Riccati equation for the covariance of estimation error. We verify the performance on simulated data sets that represent realistic human brain activities under several conditions including percentages of active sources, a number of EEG electrodes and the location of active sources. The performance of estimating brain networks is compared with a two-stage approach using source reconstruction algorithms and VAR-based Granger analysis. Our method achieved better performances than the two-stage approach under the assumptions that the true source dynamics are sparse and generated from state-space models. The method is applied to a real EEG SSVEP data set and we found that the temporal lobe played a role of a mediator of connections between temporal and occipital areas, which agreed with findings in previous studies.

## 1 Introduction

This paper aims to explore effective connectivity of underlying neural network from EEG signals. It is of great importance in neuroscience to study the direction of network connections among regions of interest (ROI) or neural nodes. Common methods of exploring directional connectivity include dynamic causal modeling (DCM), Granger causality analysis (GC), directed transfer function (DTF), partial directed coherence (PDC) that can be applied to several brain modalities such as EEG, MEG, fMRI; see a recent review in [HAVS^+^19] and detailed mathematical description of connectivity in [PS16]. In our scope, we limit ourselves to EEG analysis due to the equipment economy compared to other brain acquisitions. If only brain signals on a scalp level are available (that certainly lack of spatial resolution), we explore what more we can improve in effective connectivity analysis. This section describes literature on Granger-based brain connectivity studies examined on EEG signals. It can be categorized into two themes: one that infers brain connectivity of scalp signals and the other that concludes a connectivity in the source space. A conclusion from this survey provides us a guideline to build up our proposed model.

### 1.1 Connectivity on EEG Signals

This analysis is performed on the scalp EEG signal using a measure of dependence of interest. One typical approach is to fit a VAR model to EEG time series and use a measure such as direct transfer function (DTF) as a dependence measure in [GPO12, §4]. The sensor signals are fitted to a VAR model by least-squares estimation and then Granger causality can be obtained by performing significant tests on VAR coefficients. For example, [ACM^+^07] learned brain connectivity from VAR coefficients using DTF (directed transfer function), PDC (partial directed coherence) and direct DTF (dDTF) from high-resolution EEG data set. Moreover, a state-space framework can be applied to learn connectivity on sensor space, which is introduced in [STOS17]. The state-space model based on switching vector AR (SVAR) model was introduced for non-stationary time series, a characteristic that has been typical for biological signals. The SVAR model was represented in a state-space representation and the switching parameters were selected by a hidden Markov chain. As a result, the connectivity was learned from PDC that computed from the estimated VAR coefficients. However, it can be shown that this approach could result in *spurious causality* as mentioned in [HNMN13] where no interactions in the source level may lead to substantial interactions in the scalp level.

### 1.2 Connectivity on reconstructed sources

EEG signals cannot explain the true dynamic of neurons inside the brain because of volume conduction effects. An approach of estimating source time series from EEG signals has been developed and is referred to as *source reconstruction* or *source imaging*. The main idea is to estimate *x*(*t*) from the lead-field equation:

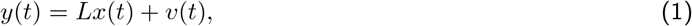

where *y*(*t*) ∈ **R**^*r*^ is the EEG data, *x*(*t*) ∈ **R**^*m*^ is the source signal, *L* ∈ **R**^*r*×*m*^ is the *lead field matrix* (given) and *v*(*t*) ∈ **R**^*r*^ is a measurement noise. The lead-field equation (1) can be used to generate artificial EEG signals when *x*(*t*) is simulated (known as *forward problem*). On the other hand, constructing the transmitted signal from the measurements in the above linear equation is often called an *inverse problem*. In order to solve the inverse problem in practice, we note that the lead field matrix varies upon several factors such as locations of EEG sensors, size or geometry of the head, regions of interest (ROIs) and the electrical conductivity of brain tissues, skull, scalp, etc. [SC13]. Examples of existing methods in source reconstruction are Low resolution tomography (LORETA), the weighted minimum-norm estimate (WMN), the minimum-current estimate, linearly constrained minimum-variance (LCMV) beamforming, sparse basis field expansions (S-FLEX) and the focal underdetermined system solution (FOCUSS) [Hau12, §2], [SC13, HNMN13, LWVS15].

In general, the number of EEG channels is lower than the number of sources. Hence, *L* is generally a fat matrix. As a result, the source imaging problem becomes an underdetermined problem. [MMARPH14] proposed that the source time series matrix is factorized into coding matrix *C* and a latent source time series *z*(*t*), then *x*(*t*) = *Cz*(*t*) where *C* is assumed to be sparse. The relationship between sources and sensors is then explained by

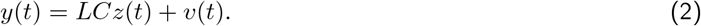

The problem of reconstructing *x* is now to estimate *z* and *C* instead. [MMARPH14] applied an *𝓁*_2,1_ regularization method by penalizing the rows of the matrix with the 2-norm to induce a sparsity pattern in source time series. Then the regularized EEG inverse problem with *𝓁*_2,1_-norm penalty term was proposed as

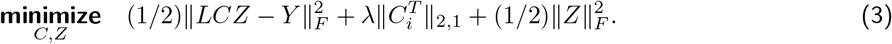

The problem is non-convex in *C* and *Z* (the matrix of latent time series.) An alternating minimization algorithm can be used for solving a bilinear problem by using initial latents *z*(0) and approximating rank of *C* from SVD. Another related approach is [WTO16] that applied sLORETA method to estimate source signals *x*. PCA was used to reduce dimension of the source signals then the principal source signals 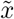 were explained 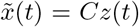,resulting in a factor model (2) and the dynamics of *z*(*t*) was explained by the VAR model. The dynamics of *x* can then be explained by the VAR model and VAR coefficients are functions of *C*.

We note that brain connectivity learned from a source reconstruction approach mainly depends on the performance of the source imaging technique. If the source reconstruction does not perform well, learning brain networks from reconstructed sources could lead to a misinterpretation.

### 1.3 Connectivity inferred from source and EEG coupled dynamics

This approach considers the dynamics of both source and sensor signals concurrently where the estimation of model parameters can infer brain connectivity directly. The work including [Hau12, HTN^+^10, GHAEC08, CWM12] considered the same dynamical model that the source signals (*x*) are explained by a VAR process and EEG signal (*y*) is a linear combination of the sources as

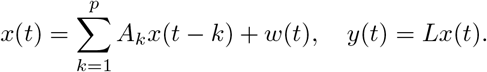

The technique to estimate unknown sources and lead field matrix (*L*) from only available mixture EEG data is called *blind source separation*. Independent component analysis (ICA) is one of blind source separation techniques that was used in [Hau12, HTN^+^10, GHAEC08]. In detail, the ICA technique relies on an assumption that the innovation term of process *w*(*t*) must be generalized as a non-Gaussian distribution. [GHAEC08] assumed that the innovation term has both sub and super-Gaussian distribution. Initially, PCA was used to reduce the dimension of EEG data with the assumption that the number of EEG channels was greater than the number of sources. Consequently, the principal EEG signals were fitted to a VAR model directly and ICA was performed on the VAR innovation term for demixing source and obtain VAR coefficients. As a result, DTF was computed from the transfer function of the source in the VAR model. However, [GHAEC08] estimated VAR parameters from the sensor signals directly, so the brain connectivity was not sparse due to the volume conduction effect. [Hau12] performed convolutive ICA (CICA) on the innovation term which was assumed to be super-Gaussian hyperbolic secant distributed for ensuring a stable solution. To obtain the sparse source connectivity, model parameters, which are *L* and *A*_*k*_’s, are estimated using the sum of *𝓁*_2_-regularized maximum-likelihood method. In addition, [Hau12, HTN^+^10, GHAEC08] assumed that the noise distribution was non-Gaussian, so the decomposition of source signals from ICA had a unique solution. [CWM12] proposed an idea to perform connectivity analysis via state-space models. The state equation was described by *generalized AR model* where the innovation process has a generalized Gaussian distribution. All state-space model parameters were obtained from maximum likelihood estimation. As a result, the relationship between sources was explained by PDC computed from estimated VAR coefficients. [CRTVV10] proposed a state-space framework for finding brain connectivity; however, the sources were assumed to be described by a VAR model. Moreover, [CRTVV10] put some prior information on the lead-field matrix where the cortical regions of interest were known. The dynamical equations are given by

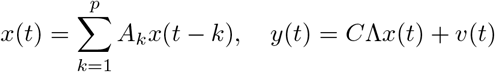

where *C* is a known matrix from a prior information on the lead field matrix and Λ is the dipole moment. When formulating the above equation into a state-space form, model parameters including *A*_1_, *…, A*_*p*_, Λ and noise covariance were estimated by expected-maximization (EM) algorithm and then Granger causality can be concluded from the estimated noise covariance. Moreover, a state-space form used in [CRTVV10, CWM12] contains source dynamics described by a VAR model and the observation equation represents a relationship between sources and sensors. The state-space parameters were estimated from maximum likelihood estimation using EM. [YYR16] proposed *a one-step state-space model* estimation framework which aims to find the connectivity in ROI level.

The state-space model used in [YYR16] was described by

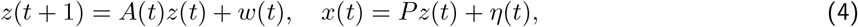

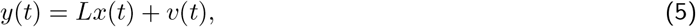

where *z*(*t*) is a time series for each ROI, *A*(*t*) is a VAR coefficient at time *t, x*(*t*) is a source time series, *P* is a binary matrix that determines sources corresponding to its ROIs and *y*(*t*) is MEG signal. Hence, the state-space model in [YYR16] is essentially a first-order time-varying VAR model to jointly explain EEG and source signals that were assumed to share a common mean activitywithin an ROI. The connectivity measure used in [YYR16] was more akin to Dynamic Causal Modeling (DCM) rather than GC, using a system dynamic matrix to explain activity flows between separate regions. The identified linear dependencies on distinct ROIs and the flow of information in cognition were represented as the product of varying VAR matrices. Nevertheless, for large ROIs, it may be unrealistic to assume that all the sources within an ROI have a common mean.

Finally, unlike other studies which used parametric approaches, [HBCN^+^17] did not assume any dynamical models relating source and EEG signals. Instead, this study was formulated around a fused lasso with a composite *𝓁*_2,1_ regularization solved numerically using the ADMM algorithm, resulting in the whole source signal being entirely and directly estimated with a sparse prior placed on some source components.

To conclude this section, learning brain connectivity from EEG data can be divided into two main approaches. The first approach explored a causality from EEG data directly (sensor space). However, a connectivity between EEG sensors is not an intrinsic connectivity explaining relationships of neuronal activities in the human brain. The second approach, consisting of *two-stage approach* and *coupled models*, was to learn brain connectivity from source signals (source space). The two-stage approach reconstructed source signals first and often explained source dynamics via VAR models. However, the performance of the two-stage approach highly depended on the performance of source reconstruction. *Coupled models* are then proposed for explaining dynamics of sources and EEG signals concurrently where brain connectivity was discovered from the estimated model parameters. Almost all previous studies assumed that source dynamics are described by a VAR process. As mentioned in [GB19] that neurophysiological data have moving-average components and should be explained by VARMA rather than pure VAR models. This paper presents a generalization of source equation to VARMA and proposes an estimation formulation based on subspace identification with a sparsity prior on the source output matrix. Contributions of this work include the following points.

▪ GC analysis of EEG signals has mostly relied on VAR models to explain source dynamics. We generalize the source equation to be a state-space (or equivalently, VARMA process), so that estimations take into account the moving-average nature of neurophysiological data. Methods of analyzing GC from estimated state-space parameters were adopted from [BS15].
▪ Practical applications of state-space GC and its theoretical properties are presented. This includes discussion of the invariant properties of GC under model coordinate transformations and signal scaling. Due to space limitation, these properties are presented in the supplementary materials.
▪ Effective connectivity will be estimated on a *moderate-dimensional* source space on the order of several ten or hundred nodes, as compared to literature [GHAEC08, HTN^+^10, CWM12, HE16a] where only a few of dipoles (≤ 10) were considered.

## 2 Background

This section describes state-space equations and the Granger characterization of this model class.

### 2.1 State-space models

Learning Granger causality of multivariate time series has been extensively applied using VAR models due to its simple linear characterization in model parameters. This paper considers a more general class of linear stochastic processes in the form of state-space equations to model EEG dynamics. We assume that source signals (*x* ∈ **R**^*m*^) is an output of state-space model whose state variable is *z* ∈ **R**^*n*^, and the EEG signal (*y* ∈ **R**^*r*^) is a linear combination of the source signals, as described in the following equations.

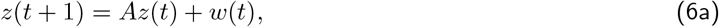

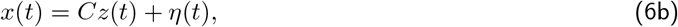

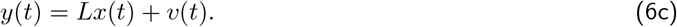

We call *A* ∈ **R**^*n*×*n*^ the dynamic matrix, *C* ∈ **R**^*m*×*n*^ an output matrix mapping the latent to source signal, and *L* ∈ **R**^*r*×*m*^ is the lead-field matrix determined from a head model. The state noise, *w*, the output noises *η, v* are zero-mean and assumed to be mutually uncorrelated.

In EEG applications, the volume conduction explains how the source signal propagates through brain tissues to the EEG signals (here from *x* to *y*) and it becomes known that Granger causality learned from *y* may not be the same pattern as one inferred from *x*, i.e., spurious effect of Granger causality [dSFK^+^16]. If model parameters (*A, C, L*) and noise covariances can be estimated from measurements *y* then we can consider (25)-(6b) and conclude a Granger causality in the source signal (*x*). In what follows, we focus on state equations of the source signal only (25)-(6b) and discuss how to learn GC of *x* once all model parameters are estimated.

### 2.2 Granger causality on state-space models

If one assumes a dynamical equation of a time series as an autoregressive (AR) process, it becomes well-known that Granger causality (GC) is encoded as a common zero pattern of all-lagged AR coefficient matrices. The generalization of this characterization to a state-space equation was provided by [BS15] and is summarized here. As our goal here is to learn a GC of the source time series, only state-space equations (25)-(6b) are considered. The noise covariance matrices in this system are *W* = **E**[*w*(*t*)*w*(*t*)^*T*^] (state noise covariance), *N* = **E**[*η*(*t*)*η*(*t*)^*T*^] (measurement noise covariance) and *S* = **E**[*w*(*t*)*η*(*t*)^*T*^] (correlation of state and measurement noise).

Granger causality concept is to determine relationships between time series from the covariance of prediction errors. If we denote 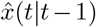 the optimal estimator of *x*(*t*) in MSE sense, it is a classical result that such optimal predictor of *x*(*t*) generated from a state-space model, based on information up to time *t* − 1 can be obtained from the Kalman filter. The Kalman filter finds the conditional mean of state variable *z*(*t*) based on all available information 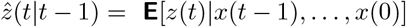and the corresponding covariance of state estimation error is 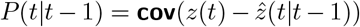. When the filter is applied in asymptotic sense, *P* converges to a steady state and satisfies discrete-time algebraic Riccati equation (DARE):

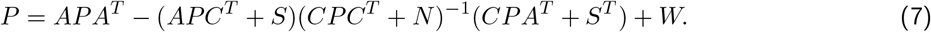

Asymptotically, the covariance of output estimation error is

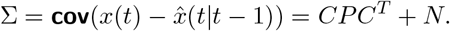

We note that if *x* ∈ **R**^*m*^ then Σ ∈ **R**^*m*×*m*^ and it is the output estimation error covariance when predicting *x* using all lagged components in *x* (full model). To determine an effect of *x*_*j*_(*t*) to *x*_*i*_(*t*) in Granger sense, we then consider the *reduced model* introduced by eliminating *x*_*j*_(*t*) from the full model, and is defined as

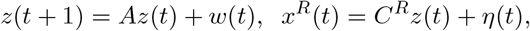

where the superscript *R* denotes the variable *x*(*t*) with *j*^th^ component eliminated and *C*^*R*^ is obtained by removing the *j*^th^ row of *C*. The optimal prediction of *x*(*t*) using all information of *x* except *x*_*j*_ is then also obtained by applying the Kalman filter to the reduced model. We can solve DARE using (*A, C*^*R*^, *W, N* ^*R*^) and obtain *P* ^*R*^, denoted as the state estimation error covariance and the output estimation error covariance of the reduced model is given by

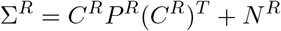

where *N* ^*R*^ is obtained from *N* by removing the *j*^th^ row and column of *N*. We also note that Σ^*R*^ has size (*m* — 1) × (*m* − 1). Doing this way, we can test if *x*_*j*_ is a Granger cause to *x*_*i*_ for all *i ≠ j* by using the Granger measure:

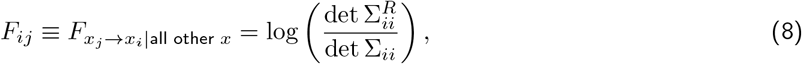

where Σ_*ii*_ and 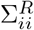are the variance of prediction error of *x*_*i*_ (*t*)obtained from using the full model and the reduced model, respectively. We can repeat the above step for *j* = 1, 2, *…, m*, i.e., learn Granger causality from data by computing *F*_*ij*_ for all (*i, j*) and construct it as a matrix whose diagonals are not in consideration. Subsequently, a significance testing is performed on the off-diagonal entries of this matrix to discard insignificant entries as zeros. We also note that the notation of *x*_*i*_ can be either a single variable or a group of variables and Σ_*ii*_ has the corresponding dimension of *x*_*i*_. When *x*_*i*_ is a single variable, then det Σ_*ii*_ reduces to the diagonal (*i, i*) entry of Σ. The resulting matrix will be called the Granger causality matrix in this paper.

## 3 Properties of GC causality

Fundamental properties of GC causality under various transformations are stated in this section. The proofs will be provided in the Appendix B.

### Theorem 1.

*For VAR process with AR coefficients, A*_1_, *A*_2_, *…, A*_p_, *we have*

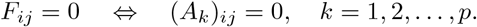

*Proof*. An example of proof can be found in [Lüt05].

It was shown in [BS15] that a Granger matrix *F* in (26) can be characterized in the state-space system matrices as well.

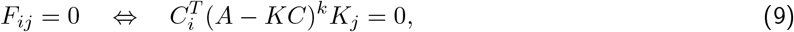

for *k* = 0, 1, *…, n* where 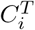 is the *i*th row of *C*, and *K*_*j*_ is the *j*th column of the Kalman gain given by

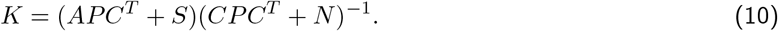

We have seen that the Granger causality condition for the VAR model is linear in AR coefficient matrices. Unlike VAR models, GC condition for state-space models is highly nonlinear in system matrices.

### Theorem 2.

*The following properties of Granger causality hold*.

1. *A GC matrix is invariant under a similarity transform of the system*.
2. *If* 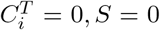 *and N is diagonal then F*_*ij*_ = 0 *and F_ji_* = 0. *As a result, the zeros of F is unchanged when N is changed under a scaling transformation*.
3. *If we permute rows of x to* 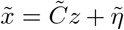, *then C is row permuted, i*.*e*., 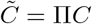 *and* 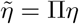. *Let* 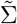 *be the covariance of*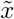. *We have* 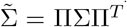. *Moreover, the GC matrix of* 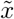 *under such permutation, is related to F by* 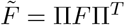.
4. *If N* = 0 *and S* = 0, *then the zero pattern of F is invariant under a scaling transformation of C*.

To interpret the meanings of Theorem 2 we consider the dynamical equations (25)-(6b) where the aim is to learn GC in variable *x*. The result in statement 1 is very natural. If one changes a coordinate system of *z*, this should not affect the causality pattern of *x*, which is the output of the linear system. For statement 2, if 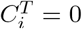, it would mean *x*_*i*_ is a pure noise *η*_*i*_. Then if *w* and *η* has no correlation, i.e., *S* = 0, and if *η* is uncorrelated, i.e., *N* is diagonal, then the effect of *x*_*i*_ cannot be transmitted to other *x*_*j*_’s by any means. Therefore, no Granger cause from *x*_*i*_ to *x*_*j*_. Similarly, *x*_*i*_ does not receive any information from other *x*_*j*_’s, so *x*_*i*_ is not Granger caused by *x*_*j*_. The invariant property of zero pattern in *F* under a scaling of *N* has a benefit when one estimates *N* in the form of *α*_*n*_*I* (homogeneous noise) and that is supposed to be a correct structure. Even when an estimated value of *α*_*n*_ can differ from the true value, the estimated zero pattern of *F* can still be correctly recovered. The statement 3 has an intuitive result. If arranging a list of brain sources in one way corresponds to a certain pattern of Granger causality then shuffling the order of brain sources (*e*.*g*., changing the brain coordinate system) leads to permuting the estimated causality pattern. Lastly, statement 4 suggests that an output signal normalization (a typical pre-processing step) can be cast as a scaling transformation of *C* under noiseless assumption. Such transformation does not change the location of null Granger causality.

## 4 Proposed method

Our proposed methodology of determining Granger causality patterns from EEG time series data consists of three main processes shown in Fig. 1, that take EEG time series and the lead-field matrix as inputs, and returns an estimated GC matrix 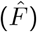 as the output. Procedure I is the core estimation formulation of the proposed EEG model using (25)-(6c) as a subspace framework. In addition to estimating the GC, procedure I provides sparsity of the estimated *C* as a by-product of source localizations. Procedure I will be explained in detail in section 4.1.

**Figure 1:**
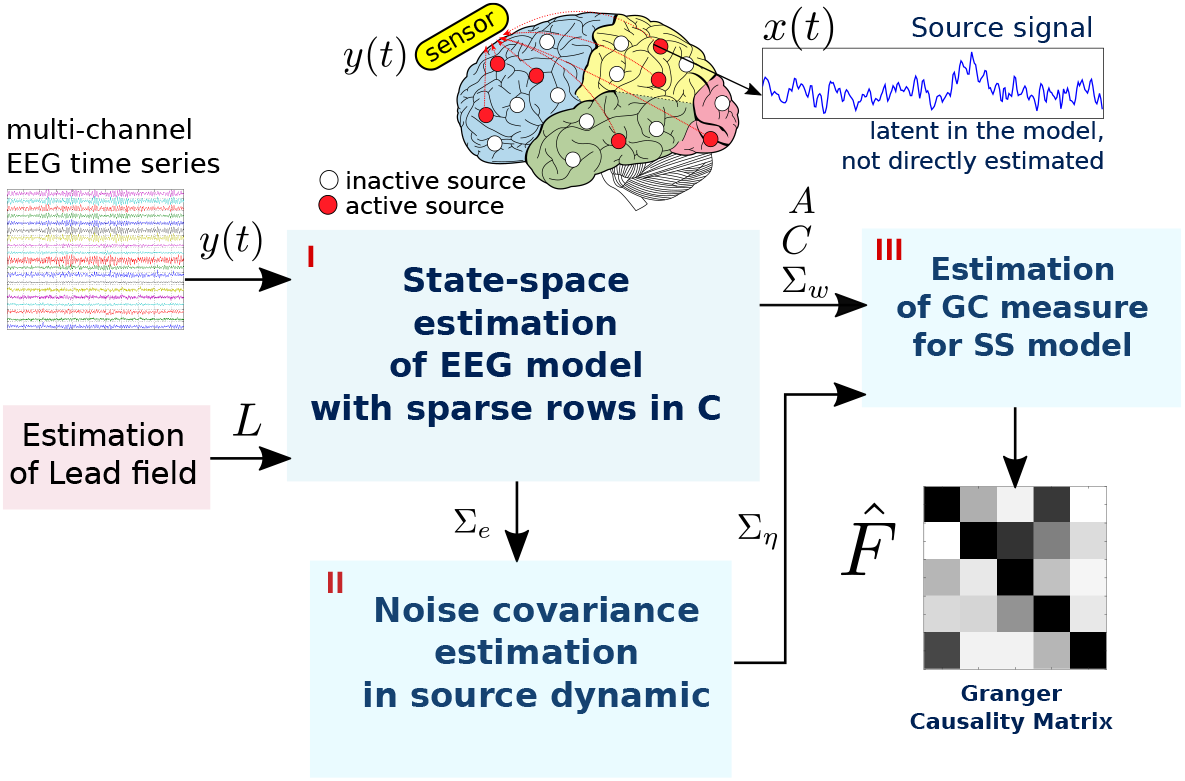
Estimation scheme for learning Granger causality from EEG data based on the proposed model.

Merging (25) and (6c), we yield

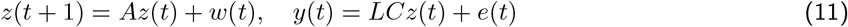

where *e* = *Lη* + *v*. The noise covariances of *w* and *e* will be employed later, represented by matrices Σ_*w*_ and Σ_*e*_, respectively. We can view *e* as a combination of noises corrupting the latent and source signals, as perceived by the output equation. After the state-space system parameters (*A, C*, Σ_*w*_) are obtained, the covariance of *e* is estimated as part of the information needed to compute the GC matrix. The noise covariance estimation, labeled as procedure II in Fig. 1 will be described in section 4.2.

The final evaluation of GC takes place in procedure III according to (26) where a heuristic threshold is used as criteria to discard small entries of *F*_*ij*_’s. Letting *F*_max_ and *F*_min_ be the maximum and nonzero minimum entries of an estimated *F*, then all values within (*F*_max_ − *F*_min_)(0, 1) in log scale that are smaller than 10^−6^(*F*_max_ — *F*_min_) will be discarded. Note that unlike the inverse algorithms or other two-stage approaches, this scheme does not estimate the source signal explicitly, but rather estimates the GC measure directly from the joint model of EEG and source signals. This approach assumes there is a mix of inactive and active sources without knowing their locations and incorporates this assumption prior to the state-space estimation.

### 4.1 State-space estimation with sparse rows in output matrix

Given the measurement data of 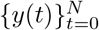, we can estimate state-space parameters *A* and *H* in (11) using the *subspace identification method* [OM12] which is available in the system identification toolbox n4sid on MATLAB. An estimated state-space model without deterministic input in this toolbox is of the form:

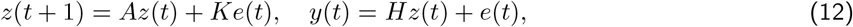

*where K* is the Kalman gain matrix and *H* in (12) takes the form *H* = *LC* according to our model (11). This section explains how to estimate *A, K, C* using a subspace identification technique with a prior structure of *C*.

Recall from (6b) that the *i*th source can be interpreted as inactive (*x*(*t*) = 0) if the *i*th row of *C* is entirely zero (in noiseless condition). To incorporate this assumption in the estimation problem, we extend the idea from our prior work [PiS18] based on a regularization technique. We put some prior in *C* by assuming that *only some sources are active* in a period of time. Consequently, *C* is *assumed to have some zero rows* corresponding to inactive sources as shown in Figure 1. We therefore propose a subspace identification framework that estimates (*A, C*) and promotes *C* to contain some zero rows.

From the subspace (stochastic) identification framework in Theorem 8 of [OM12], the main equation is

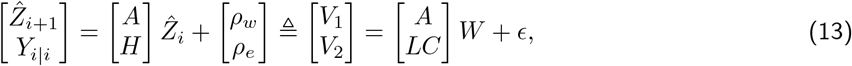

where 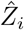 is the forward Kalman estimate of [*z*(*i*) *z*(*i* + 1)… *z*(*i* + *j* − 1)], and (*ρ*_*w*_, *ρ*_v_) are Kalman filter residuals in the innovation form (12). The key success of stochastic subspace identification is to obtain the estimated state sequence directly from the output data via an orthogonal projection. Once 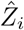 and 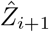 are computed, we propose to modify the existing algorithm 3 in [OM12] to estimate *C* in a regularized least-squares sense.

The algorithm of [OM12] involves the extended observability matrix

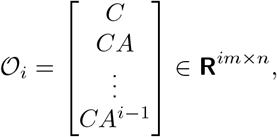

and the projection of the row space of *Y*_*f*_ (future output) on the row space of *Y*_*p*_ (past output), denoted by *ξ*_*i*_ = *Y*_*f*_ */Y*_*p*_; see complete notation details of 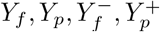 in the Appendix A. It was proved in Theorem 8 of [OM12] that the Kalman state sequences are related to the projection and the extended observability matrix via

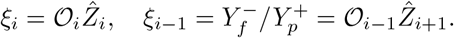

Moreover, by its definition, 𝒪 _*i*−1_ can be obtained by stripping all block rows of 𝒪 _*i*_ except the last *r* rows. From this main result, the stochastic algorithm 3 [OM12] is described as follows

1. Calculate the projections: *ξ*_*i*_ = *Y*_*f*_/*Y*_*p*_ and 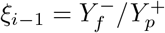
2. Calculate the SVD of the weighted projection: 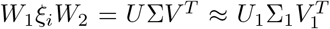 where Σ_1_ contains significantly nonzero singular values. The number of nonzero singular values determine the system order.
3. Compute 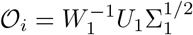 and 𝒪_*i*−1_ is obtained by extracting all rows of 𝒪_*i*_ except the last *m* rows.
4. Determine the estimated state sequences from

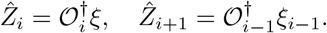

Until this step, (*A, C*) are parameters to be estimated, while other terms in (13) are known, so we propose to estimate *A* and row-sparse *C* from the following regularized least-squares problem:

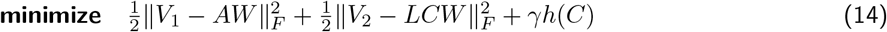

with variable *A* ∈ **R**^*n*×*n*^ and *C* ∈ **R**^*m*×*n*^ whose rows are denoted by 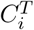 for *i* = 1, 2, *…, m*. The problem parameters are *V*_1_, *V*_2_ and *L* ∈ **R**^*r*×*m*^, the lead-field matrix computed from a head model. The estimation problem (31) is separable in *A* and *C*, so *A* is simply the least-squares solution given by *A* = (*V*_1_*W* ^T^)(*WW* ^T^)^−1^.

We propose a group-norm regularization *h* of the form

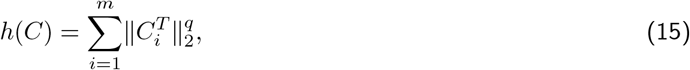

which can be regarded as a composite of *𝓁*_2_ and *𝓁*_*q*_ norms used for promoting *group* sparsity in the row of *C*. The penalty parameter *γ*, controls the degree of such sparsity, i.e., when *γ* is large, *C* tends to have more sparse rows. Choosing *q* = 1 for *h* refers to the group lasso problem [HTW15, §3.8]. In this paper, we propose to use *q* = 1*/*2 which makes *h* non-convex but this choice has been shown to obtain more desirable properties about the sparsity recovery rate [HLM^+^17] than using a convex penalty, *e*.*g*., when *q* = 1. Solving numerical solutions of the non-convex problem can be challenging. We apply a non-monotone accelerated proximal gradient (nmAPG) method [LL15] and the implementation details are explained in the Appendix C. Choosing a suitable value of *γ* in an optimal sense is a common issue in any sparse learning approach and we opt to apply model selection criterions such as BIC or AIC [HTF09]. This would require solving (31) with (15) for several values of *γ*, extracting a sparsity of rows in *C* for each *γ*, solving a constrained least-squares subject to such sparsity pattern, and selecting *γ* that yields the minimum BIC score.

We also consider the well-known *𝓁*_2_-regularization:

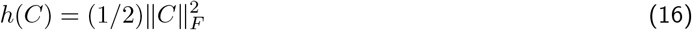

as a baseline method to compare with other estimation approaches. The *𝓁*_2_-regularized least-squares solution of *C* must satisfy the zero-gradient condition: *L*^*T*^ *LCWW* ^*T*^ + *γC* = *L*^*T*^ *V W* ^*T*^. If *WW* ^*T*^ is invertible (typically satisfied if we have enough data samples), the optimal condition can be formulated as a Sylvester equation in *C*:

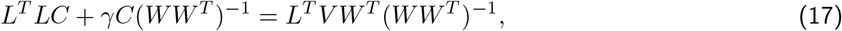

which is linear in *C* and can be solved by Bartels and Stewart algorithm (implemented in MATLAB and many linear algebra packages). The Sylvester equation has a unique solution if *L*^*T*^ *L* and − *γ*(*WW* ^*T*^)^−1^ have no common eigenvalues. Such condition holds since *L*^*T*^ *L* has nonnegative eigenvalues but− *γ*(*WW* ^*T*^)^−1^ always has negative eigenvalues [HJ13, §2]. In conclusion, the *𝓁*_2_ regularized solution of *C* is unique provided that *WW* ^*T*^ is invertible and it can be solved faster than solving (31) with the group-norm regularization. However, it is known that *𝓁*_2_-regularized solutions are not sparse. The solution *C* tends to zero only when *γ*→∞; see proof in the Appendix C. The *𝓁*_2_-regularized problem can be a remedy for a constrained least-squares of estimating *C* with a fixed sparsity pattern in rows of *C*. It is often that even *C* is constrained with some zero rows, the remaining nonzero rows still contain too many parameters, resulting in an under-determined system of solving *V*_2_ = *LCW*.

When *A* and *C* are estimated, we form the residuals *ρ*_*w*_, *ρ*_*e*_ in (13) and compute their sample covariances, denoted by Σ_*w*_ and Σ_*e*_ respectively.

#### Connection with related work

Our assumption on inferring inactive sources from sparse rows in *C* is in agreement with the use of penalty (15) by [MMARPH14]. However, [MMARPH14] did not model source dynamics but rather estimated *x*(*t*) as a whole time series segment with a prior that some components of *x*_*i*_(*t*)’s are entirely zero. For this reason, [MMARPH14] did not determine the directional GC whereas our subspace approach estimates the system parameters (not the signals) that statistically infer the causality. Elsewhere, [YYR16] employed a state-space model the same form as (25)-(6c) except that *A* was time-varying and their parameter descriptions were different from ours, namely, the state variable *z* was a mean activity in the ROI level and *C* was a binary matrix indicating whether a source belonged to a certain ROI. [YYR16]’s study did have advantage of modeling a time-varying system that not only captures non-stationarity in time series, but also provides a time-varying connectivity measure. However, such connectivity, defined from the state transition matrix, conceptually differs from GC because it explains the activity flow from one region to another, while the GC concept is based on using the information of one variable to better predict another. Moreover, [YYR16] applied an EM algorithm in the estimation process due to the presence of latent variables, while our current approach handles this issue by introducing a regularized subspace identification.

### 4.2 Estimation of noise covariance in the source dynamic

The GC estimation from state-space model parameters explained in Section 2.2 requires information of noise covariances (both state and measurement noises). Consider our methodology in the diagram 1 and the model equations (25) and (6b). At this step, we have estimated *A*, Σ_*w*_, *C* from subspace identification. Then it is left to estimate Σ_η_ (the measurement noise covariance at the source equation) in order to solve a GC matrix via the Riccati equation.

The measurement noise observed at the output equation (11) has the covariance related to the covariances of *η, v* by

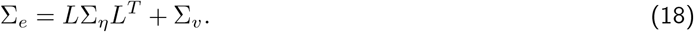

In other words, the RHS of (18) is the model-produced structure form of Σ_*e*_ where its value is obtained empirically from the subspace identification described in Section 4.1. The lead field matrix can be obtained from a head model (as part of our assumptions). Therefore, it remains to estimate the unknown Σ_*η*_ and Σ_*v*_. Consider the dimensions of all these matrices, where they are symmetric and positive definite, i.e., Σ_*e*_ ∈ **S**^*R*^ and Σ_*η*_ ∈ **S**^*m*^ and Σ ∈ **S**^*R*^. Linear equation (18) may have many solutions, so we propose to estimate Σ_*η*_, Σ_*v*_ with a certain structure in an optimal sense using the KL divergence distance with a Gaussian assumption.

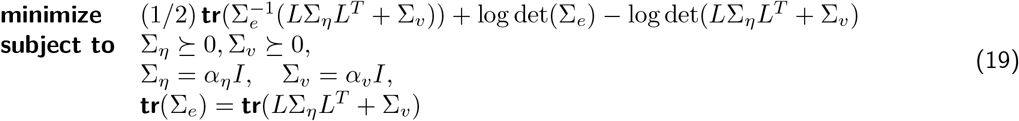

with variables Σ_*η*_ ∈ **S**^*m*^ and Σ_*v*_∈ **S**^*R*^. We choose to restrict down to a diagonal structure on the variables, corresponding to the assumption that each of the noise vectors *η* and *v* is mutually uncorrelated and has a uniform variance (*α*_*η*_, *α*_*v*_). In addition, the trace constraint in (19) explains the conservation of the noise average power, i.e., the empirical average power and that of the model-produced form must be equal.

The problem (19) can be further simplified since the variables are merely scalars of *α*_*η*_ and *α*_*v*_. The trace constraint in (19) gives the linear relation:

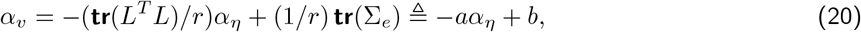

which makes the cost objective in (19) reduced to a function of *α*_*η*_ only. Moreover, the relation between *α*_*η*_ and *α*_*v*_ and the positive constraint on *α*_*v*_ results in the inequality: 0 ≤ *α*_*η*_ ≤ *b/a*. As a result, we can reformulate (19) into a *scalar* optimization problem as

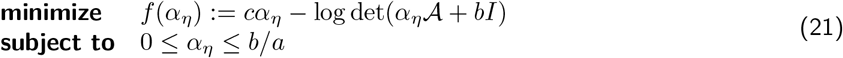

with variable *α*_*η*_ ∈ **R** and the problem parameters are 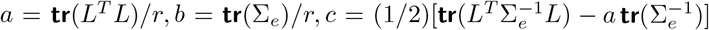, and = *LL*^*T*^− *aI*. The cost objective of (39) are convex in the variables. In fact, we describe in the Appendix E that solutions of (39) can be obtained almost in a closed-form expression depending on three-case conditions of the problem parameters and the three-case solutions are i) Σ_*η*_ = 0, ii) Σ_*η*_ has the same average power as Σ_*e*_ and iii) the noise power of *e* is decomposed to Σ_*η*_ and Σ_*v*_ in an optimal trade-off according to the optimal KL divergence.

If Σ_*e*_ ⪰ 0 (degenerated case), then KL divergence is not valid. We estimate Σ_*η*_ and Σ_*v*_ in a least-squares sense instead. That is, we minimize 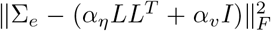 over (*α*_*η*_, *α*_*v*_) and the covariance estimates are Σ_*η*_ = *α*_*η*_*I*, Σ_*v*_ = *α*_*v*_*I*.

### 4.3 Learning significant Granger causality

From the scheme proposed in Figure 1, after we have estimated a Granger causality matrix, 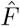, one needs to decide which (*i, j*) entries of *F* are significantly (or statistically) nonzero. Statistical tests on GC measures characterized from VAR models are available as the log-likelihood ratio test for a nested VAR model or GC inference tests on autocovariance sequence and cross-power spectral density are provided in a MATLAB toolbox by [BS14]. For Granger causality characterized on state-space models, [BS15] concluded that the inference measure in (26) does not have a theoretical asymptotic distribution, while it was observed in their experiments that the test statistics can be well-approximated by a Γ distribution.

As an alternative, a significance testing can be performed using permutation tests or bootstrapping methods. To perform such tests under the null hypothesis that *x*_*j*_ does not Granger cause *x*_*i*_, it often requires shuffling temporal segments of *x*_*j*_ to examine whether such randomization does not change the effect of *x*_*j*_ to *x*_*i*_. We note that the permutation is impractical to apply in our context since our proposed scheme does not estimate the source signal *x* directly. Alternatively, *Kappa selection* approach [SWF13] is a scheme used in variable selection problem to tune problem parameters (here in our context, a threshold to regard *F*_*ij*_ as zero) by a score criterion called *Kappa score*. This method also requires segmenting EEG time series; each of which is used to estimate GC matrix. The work in [PiS19] considered the vectorized version of estimated GC matrices that contain both null and causal entries, and then applied Gaussian mixture models to cluster *F*_*ij*_’s where the group having the least mean was regarded as null GC. The approach in [PiS19] relies on the central limit theorem to conclude that an averaged GC matrix estimated from multi-trial data converges to a Gaussian distribution. A limitation of approaches in this direction is a requirement of repetition of estimation processes on segmented or multi-trial data and hence, a computation power becomes a trade-off.

To the best of our knowledge, a statistical significance test of state-space Granger causality is still an open question where a challenge is on deriving the asymptotic null sampling distribution of the estimator [GB19]. We are aware of the importance of significance testing; however, this paper is not aimed to pursue this topic as it is beyond the paper scope. We will use a heuristic thresholding on discarding small entries of *F*_*ij*_’s. Let *F*_max_ and *F*_min_ be maximum and nonzero minimum entries of estimated *F*. A threshold is varied in (*F*_max_ − *F*_min_)(0, 1) in log scale where the selected threshold is 10^−6^(*F*_max_ − *F*_min_).

### 4.4 Generating EEG data

This section describes an approach for generating state-space models with some sparse GC pattern so that we can evaluate the accuracy of estimated GC with the ground-truth one. The procedure relies on an important result by [BS15, BS11]. The GC causality of a filtered VAR process is invariant if the filter is diagonal, stable and minimum-phase. Let 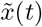 be a *p*-lagged VAR process where the *z* transform relation is given by 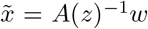 with VAR polynomial:

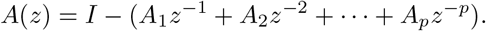

We consider *G*(*z*) an MIMO (multi-input multi-output) transfer function of the form:

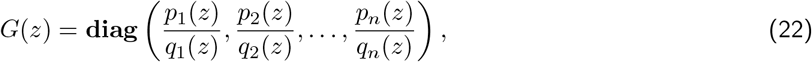

where each of diagonal entries of *G* is a rational proper transfer function of a given relative degree. The minimum-phase and stability properties of *G* suggest that the roots of *p*_*i*_(*z*) and *q*_*i*_(*z*) must lie inside the unit circle, respectively. As a result, we define 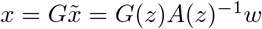 *w* and *x* is a VARMA process. The result from [BS11] shows that *x* also has the same GC pattern as 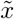, which is easily explained from a zero pattern in VAR coefficients. The system transfer function from *w* to *x* can be equivalently represented in a state-space form. Therefore, we proposed a procedure to generate a state-space equation with sparse GC pattern as follows.

1. Generate sparse *A*_1_, *A*_2_, *…, A*_*p*_ matrices randomly with a common zero pattern and the polynomial *A*(*z*) must be stable. This is to guarantee that the generated VAR process is stationary. We can do this by randomizing stable roots inside the unit circle and compose the polynomial in the diagonal of *A*(*z*). Consequently, off-diagonal entries of *A*_k_’s are generated randomly in a common (*i, j*) location. If the resulting *A*(*z*) is not stable, we randomize off-diagonal entries again. In practice, when *n* is large (in order of several tens or hundred), it is getting more difficult to obtain stable VAR unless the VAR coefficients should be very sparse.
2. Generate a random diagonal transfer function *G*(*z*) with required properties. We can generate stable zeros and poles of *G*(*z*) when the orders of two polynomials are given.
3. The transfer function from *w* to *x*, the desired source signal, is then given by *H*(*z*) = *G*(*z*)*A*(*z*)^−1^. Convert *H* into a discrete-time state-space form using `tf2ss` command in MATLAB. We obtain (*A, B, C, D*) of the state-equation: *z*(*t* + 1) = *Az*(*t*) + *Bw*(*t*), *x*(*t*) = *Cz*(*t*) + *Dw*(*t*). Since *H* is a proper transfer function, we have *D* = 0.

State-space equations and VARMA models can be interchangeably transformed [CGHJ12], so we can refer to the generated model as state-space or VARMA model with sparse GC pattern.

**Special case of** *G*(*z*). As suggested in [BS15] is when *G*(*z*) in (22) has the form of a minimum-phase MA polynomial: *G*(*z*) = (1 + *cz*^−1^)^q^*I* = *C*(*z*)*I* with |*c*| *<* 1, the model reduces to *x* = *G*(*z*)*A*(*z*)^−1^*w* = *C*(*z*)*A*(*z*)^−1^*w* = *A*(*z*)^−1^*C*(*z*)*w* since *C*(*z*) is just a scalar. We can then readily consider *x* as a VARMA(*p, q*) process. The AR and MA coefficients in *A*(*z*) and *C*(*z*) can be used to convert into a state-space form, for example, the Hamilton form.

As described above, we have generated parameters of ground-truth models of source signals according to (25) and (6b). It remains to generate a lead field matrix, *L*, which is computed based on the New York head model described in [HPH16] and select model parameters corresponding to realistic assumptions on EEG signals. We follow implementation details in [HE16a] and add some extensions: i) more number of sources can be considered *m >* 2, ii) source dynamics are VARMA (not VAR) and iii) source time series have an underlying Granger causality, which is generated randomly.

## 5 Performance evaluation

We aim to evaluate the performance of our method on simulated EEG data sets first. In the data generating process, we can set up model dimensions (*n, m, r*) and a ground-truth sparsity pattern on the GC matrix associated with such models. In an estimation process, one needs to assume the model dimension; here let 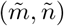 be the number of sources and latents in the estimation which could be larger or smaller than (*m, n*), while *r* (the number of EEG channels) is certainly known. Then it leads to a condition in an evaluation procedure since the estimated matrix *F* of size 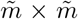 could have a different dimension from the ground-truth matrix *F*. Recall that a ground-truth model used to generate data is explained in (25)-(6c). We describe how to calculate the classification measures in a fair setting. In this study, we assume that 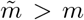 since we can overestimate the number of sources and we expect the source selection procedure to remove inactive sources at the end. By this assumption, 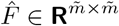 has a bigger dimension than the true Granger causality matrix *F* ∈ **R**^*m*×*m*^.

Figure 2 shows all three square regions involved in the evaluation process. We start with the **true source region (T)** that contains all the sources in a ground-truth model, and since not all sources are active, a subset called **active source region (A)** consists of all the true active sources where we can reorder the source coordinates so that active sources contain in this region. We define the **estimated source region (E)** as the set of all sources considered in an estimated model. By the assumption that 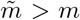, then the true source region must lie inside the estimated source region. By these notations, the set *T* − *A* contains all inactive sources in the ground-truth model (highlighted in the blue color), and *E* − *T* (green area) represents possible Granger causality that occurred in estimated sources that do not exist in the ground-truth model. The circles ° denotes the predicted nonzero GC 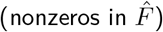, and the black circles are true positive (TP) and while the red circles are false positive (FP). The cross signs denote the predicted zero GC 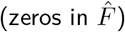, and the red crosses are false negative (FN) while the black crosses are true negative (TN). Hence, when we evaluate an estimated GC matrix 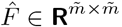 the following properties hold on the regions shown in Figure 2.

**Figure 2:**
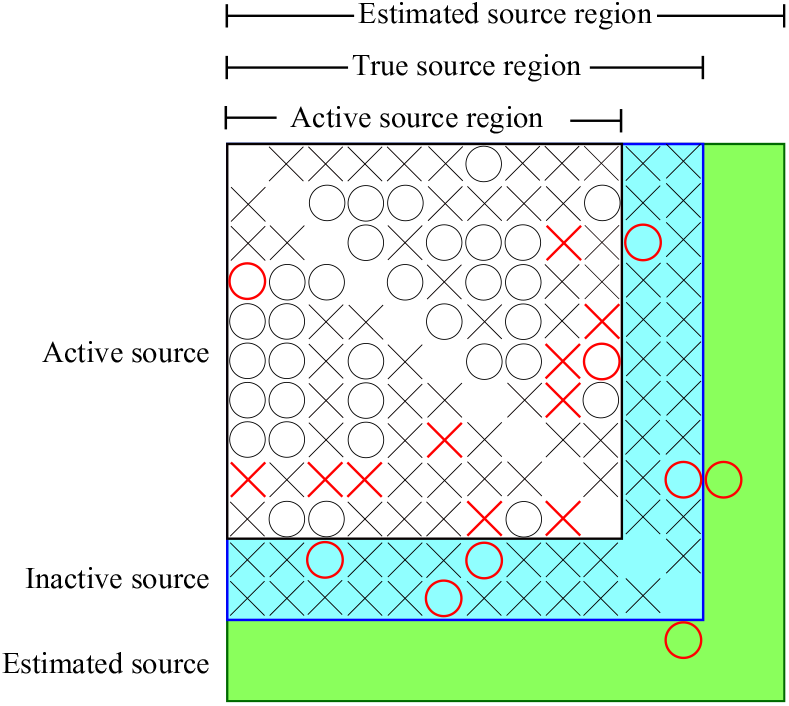
Granger causality evaluation including active source regions, true source region and estimated source region.

▪ TP and FN only exist inside the active regions because nonzeros of *F* in a ground-truth model can only exist in this region.
▪ True positive rate (TPR) is equal in all regions because the numbers of TP are equal in all regions.
▪ If all active sources are correctly classified then there is no FP in the true source region and the estimated source region.
▪ Predicted nonzeros in the green region are regarded as FP since there are no true sources there.
▪ A fair comparison should be tested on the true source region.
▪ Accuracy (ACC) and True negative rate (TNR) between regions cannot be compared because the numbers of negatives are different in those regions.
▪ FP and FN on the estimated source region can only be evaluated when a method is tested on a simulated data sets as the ground-truth models and hence the true source region are known.

From above reasons, the performance on the active true source region reflects how well the method can achieve in TPR. An overall performance of a method can be worse when evaluated on the true source region since if the method predicts any nonzero in the inactive source region, it must be FP. A good method should yield a high TNR on the blue area. Lastly, the performance evaluated on the estimated source region can only drop if the method introduces unnecessary predicted nonzeros in the green area. This arises from two possibilities: error from the source selection algorithm or error from learning significant GC entries.

## 6 Simulation results

The number of state variables (or latents), sources, and EEG channels in the ground-truth models are denoted by *n, m, r*, respectively. In the model estimation process, *m* and *n* must be set and we use a notation of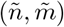 (which are not necessarily equal to (*n, m*). Therefore, 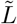 also denotes the lead-field matrix calculated from the parameter 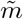. Three main factors to the performance of active source selection and estimating GC causality are as follows.

1. The percentages of active sources in the ground-truth model are set to 20% and 40% with *m* = 50.
2. The number of EEG electrodes varies as *r* = 108, 61, 31, 19.
3. The percentages of deep sources in the ground-truth model varies as 0%, 50%, 75%.

According to [HE16b], we define eight regions of interest (ROI) that cover left-right, anterior-posterior, and superior-inferior hemispheres, and are labeled as RAI, RAS, RPI, RPS, LAI, LAS, LPI and LPS. Ground-truth models are assumed to contain 8 ROIs and all sources (including both active and inactive) are drawn from 4 ROIs randomly. The first two factors are considered to examine how the performance depends on the sparsity of the ground-truth system and the number of measurements. The third factor is known to affect a performance of localizing active sources. The fourth factor, we aim to investigate the robustness of the method when we could wrongly choose the model order in the estimation 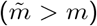,which is a common aspect in practice. As we vary the above three factors, we obtain 2 × 4 × 3 = 24 cases to show performances of our source selection and Granger causality learning approach.

For each fixed (*n, m, r*) and each controlled factor, we randomly generated 100 ground-truth models with different underlying GC causalities and corresponding 100 realizations of EEG time series with SNR of 0.95. All classification performance indices: true positive rate, false positive rate, accuracy, F1 score (TPR, FPR, ACC,F1) are averaged over 100 runs. The ground-truth VARMA models are generated with the sparse VAR part of dimension: 10, 20, lag of order 2, and the diagonal filter (moving average part) of order 6.

There are many source reconstruction algorithms that can be compared with our source selection scheme (31). Moreover, Granger causality can be estimated from the reconstructed sources from these inverse algorithms using VAR-based approach, as implemented in MVGC toolbox [BS14], and then compared with our estimated state-space Granger causality. For this purpose, we also implemented source reconstruction algorithms including the weighted minimum norm estimator (WMNE), the linear constraint minimum variance beamformer (LCMV), and the standardized low-resolution brain electromagnetic tomography (sLORETA); all implemented in Brainstorm [TBM^+^11], freely available for online download under the GNU general public license (http://neuroimage.usc.edu/brainstorm). In WMNE implementation, the depth weighting is set to 0.5; the regularization parameter of the noise method is set to 0.1 and SNR is fixed as 3. As for LCMV, the noise covariance regularization uses the median value of the eigenvalues. In sLORETA, no depth weighting is applied and SNR is fixed as 3. In the comparative experiments with Brainstorm, we set (*m, r*) = (50, 61) with 20% active sources and 50% deep sources in ground-truth processes. In our estimation process, we set 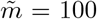 and the assumed potential sources are scattered over 8 ROIs of the brain.

All simulation data sets and codes used in this paper are available at https://github.com/parinthorn/eeg-bc.

### 6.1 Selecting active sources

In this experiment we show the performance of classifying active sources. Figure 3 shows a typical example of estimated *C* when the number of all sources in estimation could be falsely larger than the actual number 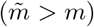. The estimated source index *i* when *i > m* is regarded as spurious active source if it is incorrectly detected as an active one by inferring from nonzero rows of 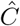. Our method returned a very small percentage spurious sources in Figure 3 showing a capability of the *𝓁*_2,1/2_ regularized estimation to select sparse rows in *C* with a good accuracy.

**Figure 3:**
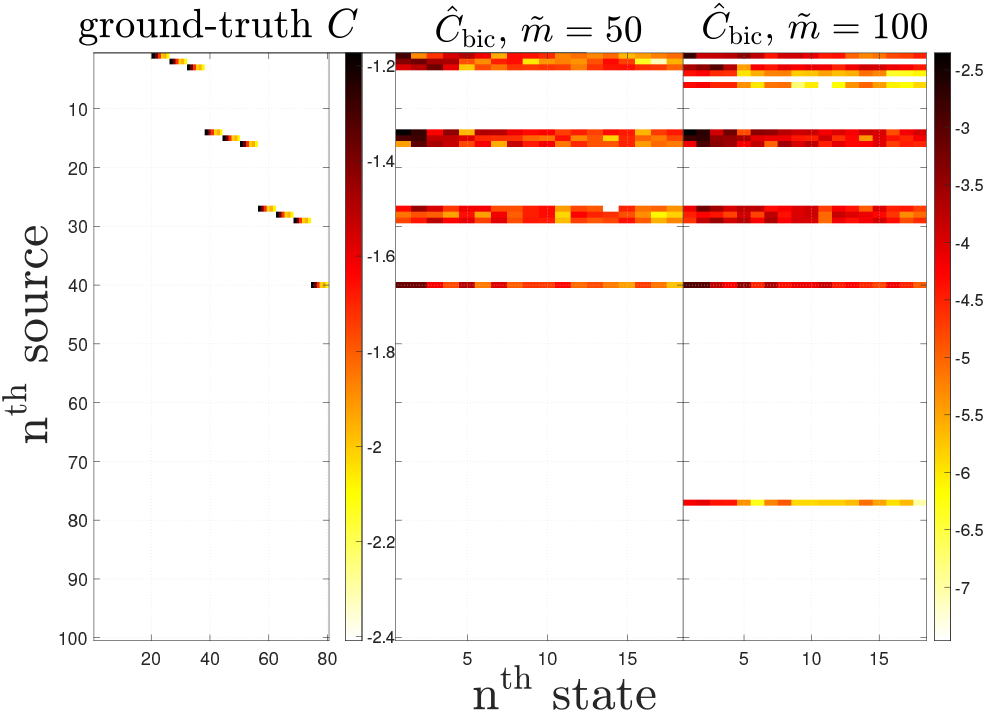
Examples of zero patterns of estimated *C* as 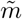 varies. The color scale is proportional to magnitudes of *C*_*ij*_’s in log scale. The regularized solution of 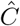 corresponds to the use of *λ* chosen from BIC. The percentage of deep source is 50% and the number of electrode is 61.

Classification performance of the method which depends on *λ* in the formulation (31) is shown in Figure 4. Each point on ROC curves refers to a classification result from a value of *λ*, where true positives (negatives) correspond to correctly identified nonzero (zero) rows in 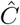. The ROC plots show that the performance of classification varies upon the number of EEG channels. As we have more electrode measurements (more data samples), the ROC curve is shifted toward the top left corner (improved accuracy). The results confirm with [SYW^+^16] that at least more than 64 electrodes are needed for source reconstructions with good quality. Figures 4 (a) show superior performances if the number of active sources is relatively small because a sparsity-inducing formulation (31) generally works well when ground-truth models are sufficiently sparse [HTW15]. Our result in Figure 5 also shows the main factor to source selection performance, which is the location of active sources. As the ratio of deep sources to shallow sources is higher, the smaller area under the ROC curve is obtained.

**Figure 4:**
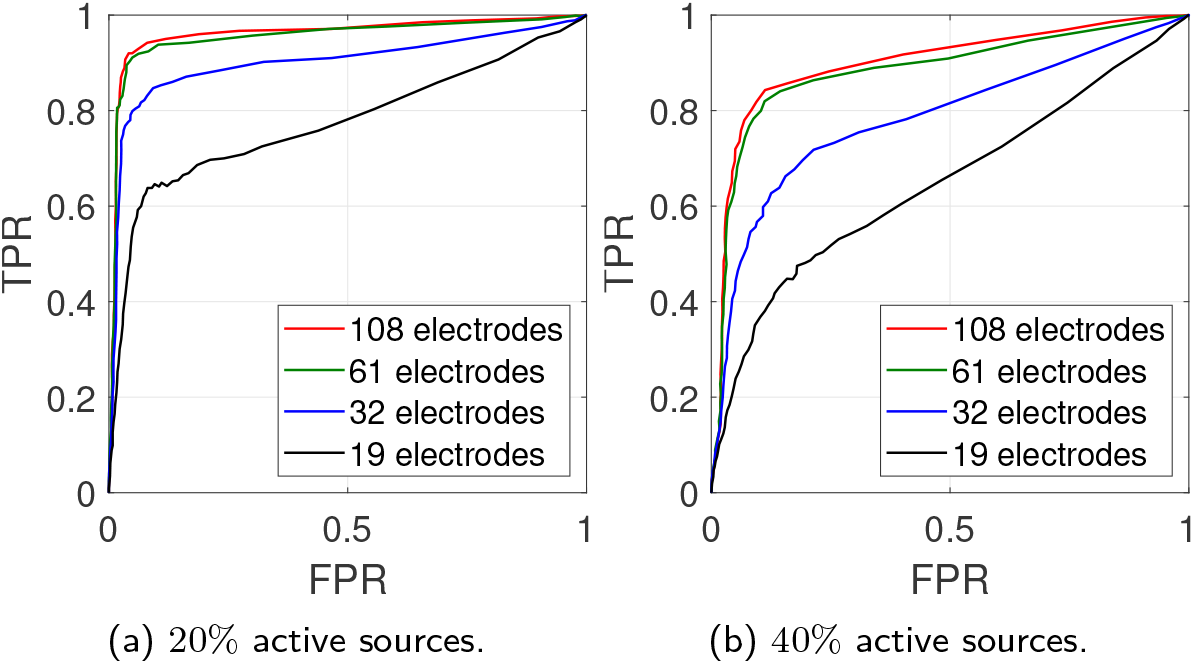
Receiver operation characteristic (ROC) of active and inactive source classification as the number of EEG sensors varies.

**Figure 5:**
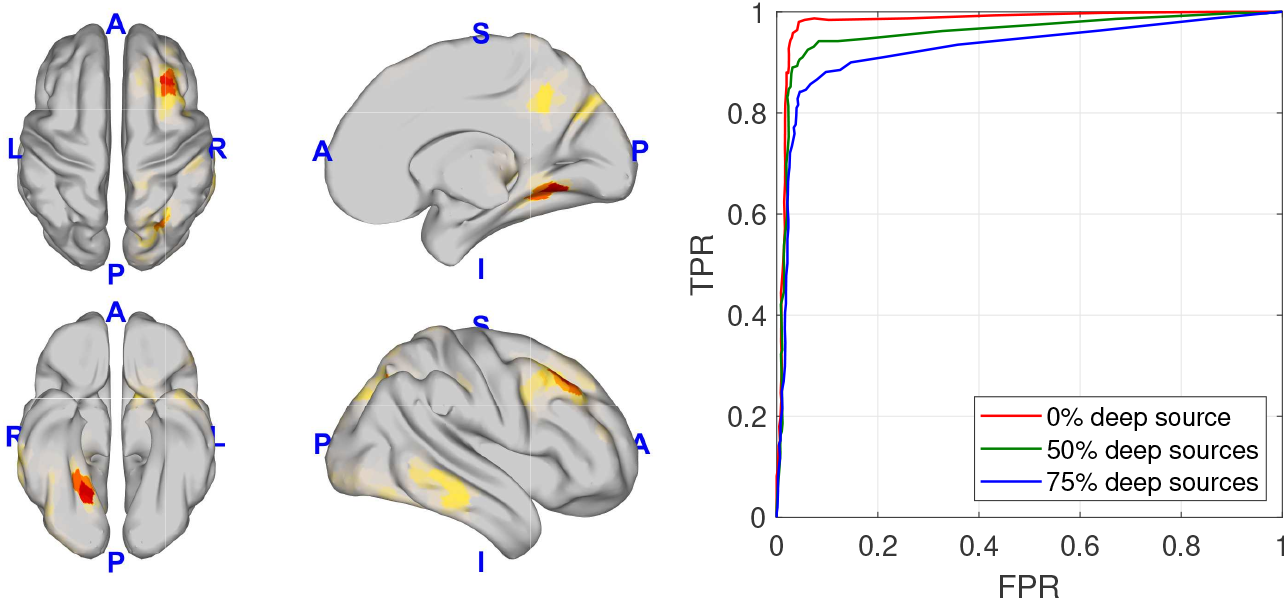
*Left*. An example of ground-truth source locations. *Right*. Receiver operating characteristic (ROC) of active and inactive source classification as varying the percentage of deep sources. The percentage of active sources is 20% and the number of electrodes is 61.

ROC curves only explain how classification performances vary under the parameter *λ*. In this experiment, we evaluated the source selection performance when *λ* was chosen by BIC and displayed it by box plots of TPR, FPR, and ACC in Figure 6 as the distribution of these 100-run metrics may be skewed. TPRs of higher than 80% were mostly obtained when using 61 or 108 electrodes and given that the number of active source is small. Our approach has a great advantage in achieving almost zero FPR when the percentage of active sources is small as seen in Figure 6 (left) that the median of FPR almost goes to zero. When the ground-truth sources are more active, more portions of higher FPRs and TPRs decrease to under 80%. The overall accuracy in the case of 20% active sources is not sensitive much to the number of electrodes, as compared to the case of 40% active sources.

**Figure 6:**
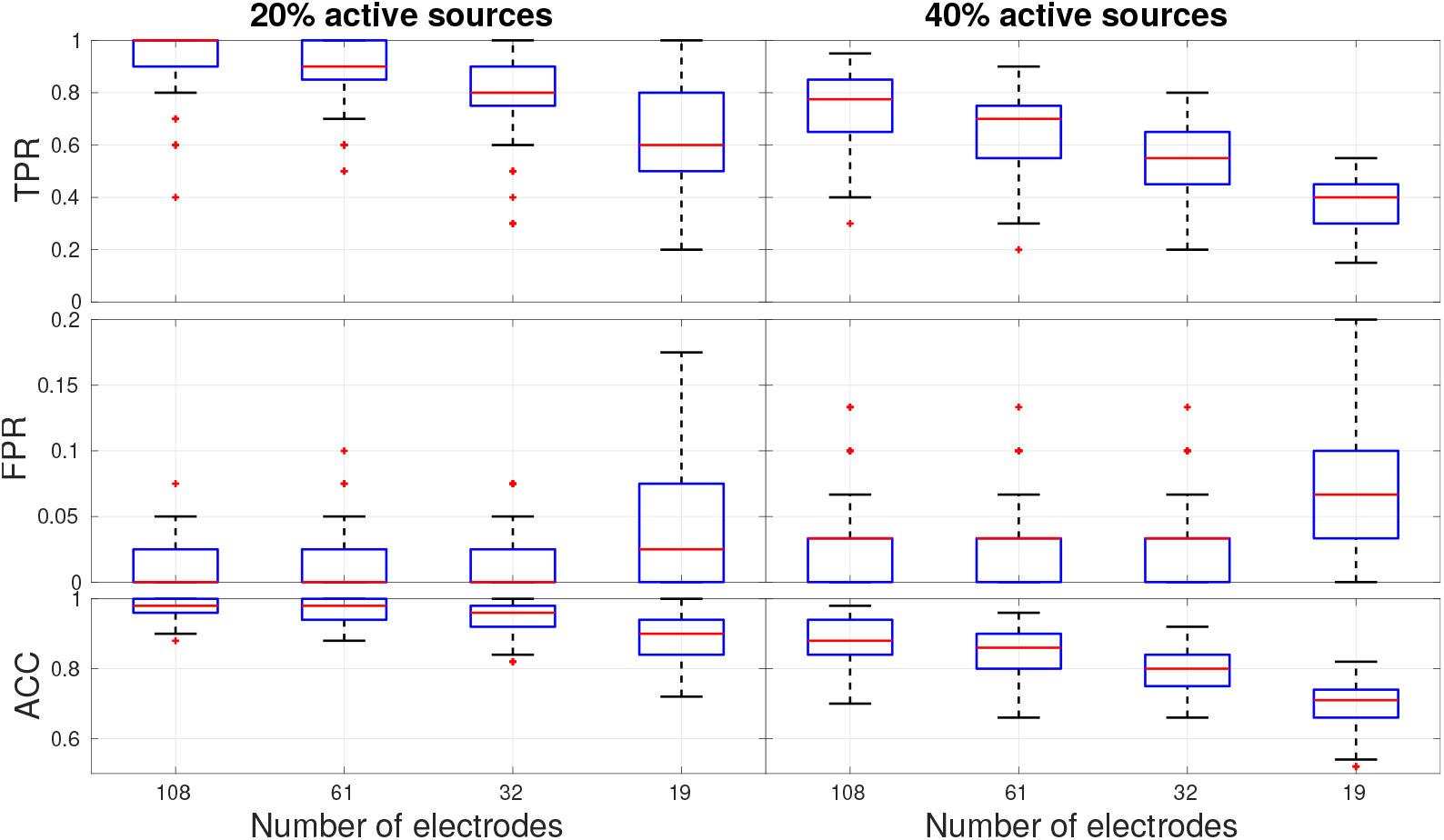
Box plots of source selection performance metrics (TPR, FPR, Accuracy) as the number of electrodes varies and under two conditions of the number of true active sources.

Figure 7 compares source selection results with WMNE, LCMV, and sLORETA. An example from the 25th run out of all 100 samples available in https://bit.ly/3jAJEeS, shows that the true active sources can be typically recovered by our method. For source reconstruction algorithms, the ground-truth deep sources in the left hemisphere can be mostly detected by LCMV but not by WMNE and sLORETA. This agrees with a comparison of inverse algorithms given by [APS^+^19]. It was concluded that when source locations are deep, the overall accuracy of LCMV is higher than eLORETA (which is an improved version of sLORETA) in a high SNR setting (which is the case in this experiment).

**Figure 7:**
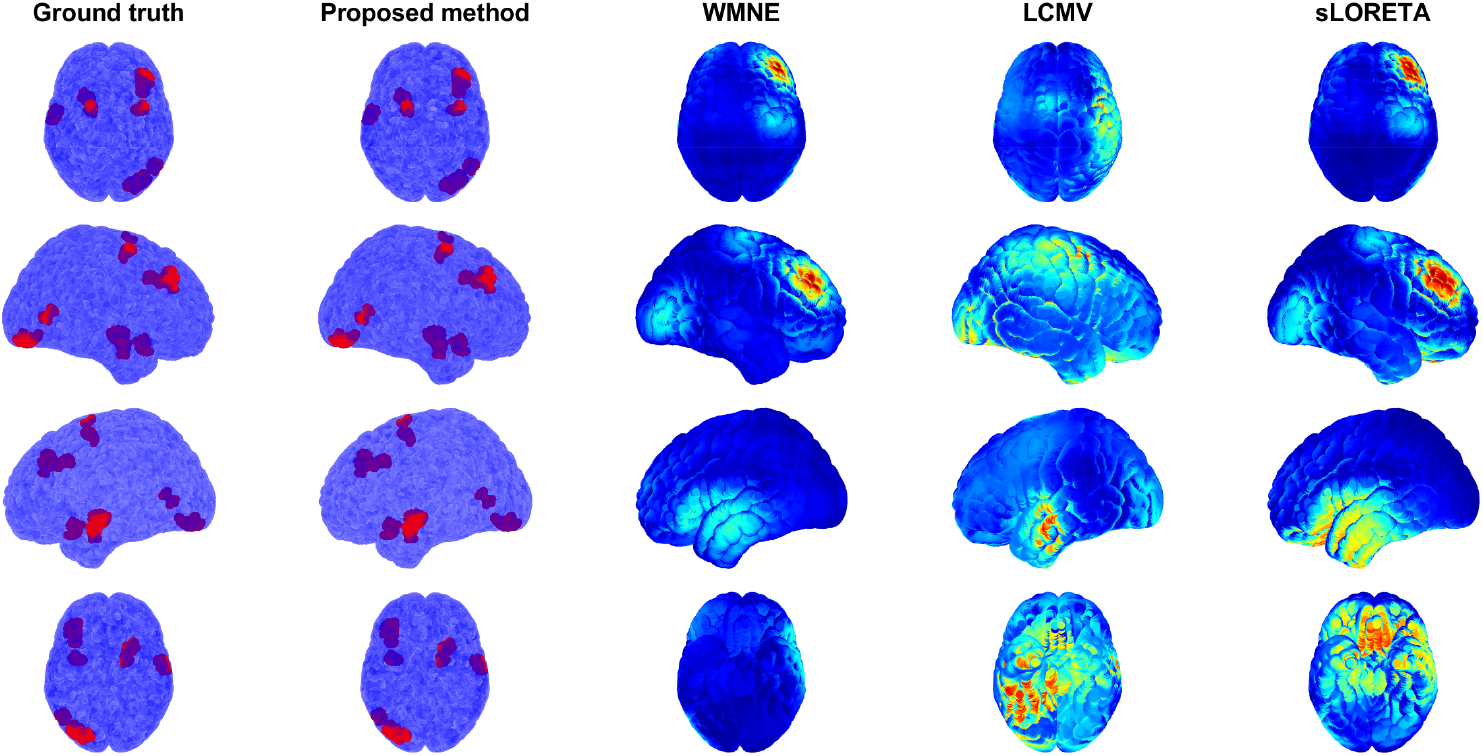
Comparisons with source reconstruction methods: WMNE, LCMV, sLORETA. Our averaged performance indices over 100 runs (of recovering 50 sources) are (TPR,FPR,ACC) = (0.8130, 0.0172, 0.9658).

### 6.2 Estimation of Granger causality

The results of discarding inactive sources in section 6.1 showed that if a ground-truth system contains only a few active sources, our method can select the active ones with a good accuracy. In this experiment, we explore Granger causality pattern among the selected active sources. Hence, we show the performance of estimating GC as a binary classification problem (regard *F*_*ij*_ as null or causal entry) from simulated EEG signals that were described in section 6. The performance indices, TPR, FPR, ACC and F1 score are reported as three factors (sparsity of ground-truth, percentage of deep sources, and the number of EEG channels) vary.

Our performance of estimating GC was compared to a two-stage approach where a VAR-based GC was learned from reconstructed sources estimated by the inverse algorithms (WMNE, LCMV, sLORETA). We showed this result in a setting that i) the total 50 sources contained 20% active ones, ii) 50% sources were located in deep ROIs and iii) the number of electrodes was 61. In Brainstorm implementation, reconstructed sources in the resolution of 2000K were averaged within each area of 8 ROIs. For GC estimation, the MVGC toolbox [BS14] was implemented using autocovariance method (LWR); VAR lag orders were selected from BIC; significance level of testing VAR-based GC is set to *α* = 0.01. For our method, we set 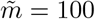 and sampled these 100 potential sources over 8 ROIs. The method can return an estimated GC matrix in node-based resolution 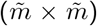or ROI-based resolution (8 ×8) depending on how we cluster a group of variables (*x*_*i*_, *x*_*j*_) in (26).

Figure 8 shows that our method generally performs well in all metrics when the ground-truth source dynamics contain fewer active sources. TPR was higher when the number of electrodes was reduced since we had less data samples. As our formulation and the scheme of selecting the regularization parameter based on BIC favor sparse models, FPR and ACC had small variations as the number of electrodes changes. In overall, the accuracy of sparse ground-truth case was above 95% and not sensitive to the number of electrodes. In the case of denser ground-truth models, we observed a different trend on FPRs, contrary to the case of 20% active sources. As we use fewer EEG channels, data used in estimation were less and BIC tended to select fewer active sources. As stated in (28), the selected zero rows in *C* always infer zero rows and columns in the GC matrix, i.e., inactive sources have no GC with any other sources. This implied a decreasing trend of FPRs as the number of electrodes is less. The overall accuracy then had slightly increasing variations in this case. If we focus on the performance of correctly classifying causal entries of a GC matrix, out of all predicted nonzeros, then F1 scores were obtained in the range of 10 − 35%, and they were decreasing as the number of electrodes is less. The overall accuracy was above 90% due to the sparsity-promoted framework in the source selection that allowed us to predict the null entries of GC correctly.

**Figure 8:**
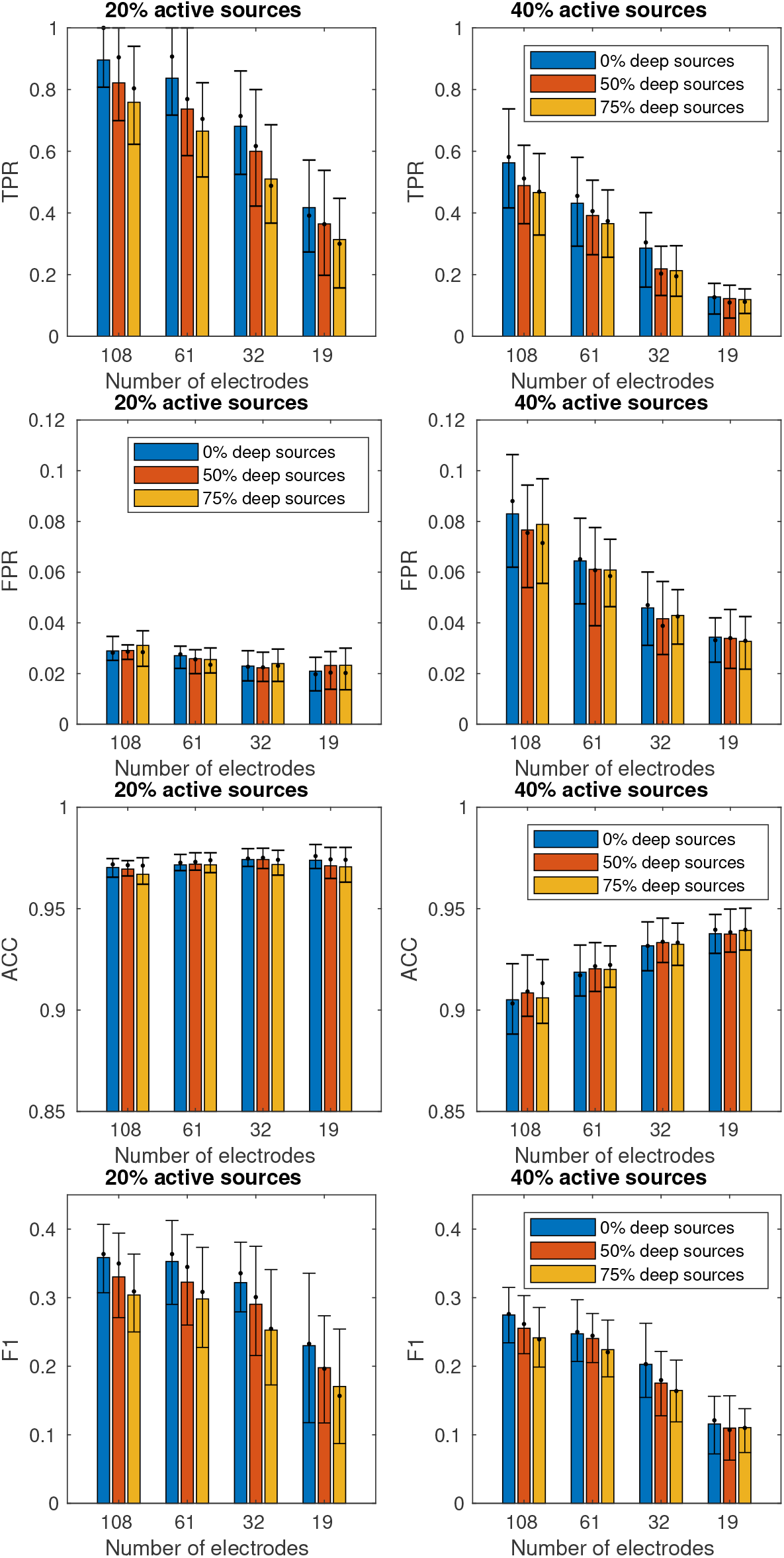
Performance indices of classifying GC causality. The color bar is the averaged performance over 100 runs. The dot is the median and the vertical bar represents the interquartile.

We showed a typical example of estimated GC from the 25th run which is mapped in ROI-based resolution in Figure 9 and the averaged performance over 100 runs in Table 1. The ability to rule out inactive sources in our method helped improve detecting zero entries in the GC matrix. This led to a significant reduction of false positives as compared to WMNE, LCMV, and sLORETA. Whereas LCMV can relatively recover more deep sources than other inverse algorithms, the inherent moving average dynamics of the ground-truth sources had made it difficult for the VAR-based GC estimation method to accurately recover GC from the reconstructed sources. As we observed from Fig. 9, there were biases in strong GC connections inferred from VAR estimates via the two-stage approach. The unexplained moving average dynamic in the residual of estimated source signals can introduce errors in VAR estimates and hence in false GC connections. This was supported by the simulation that while we have set the lag order of the VAR part to 2 in the ground-truth system, the VAR estimation typically picked higher order around 6. In contrast to the two-stage approach, our framework provided a means for estimating VARMA parameters directly in a state-space form. In combination with a prior on sparse rows of *C*, our method favored sparse GC networks, which further significantly improved overall accuracy, provided that the ground-truth network is also sparse.

**Figure 9:**
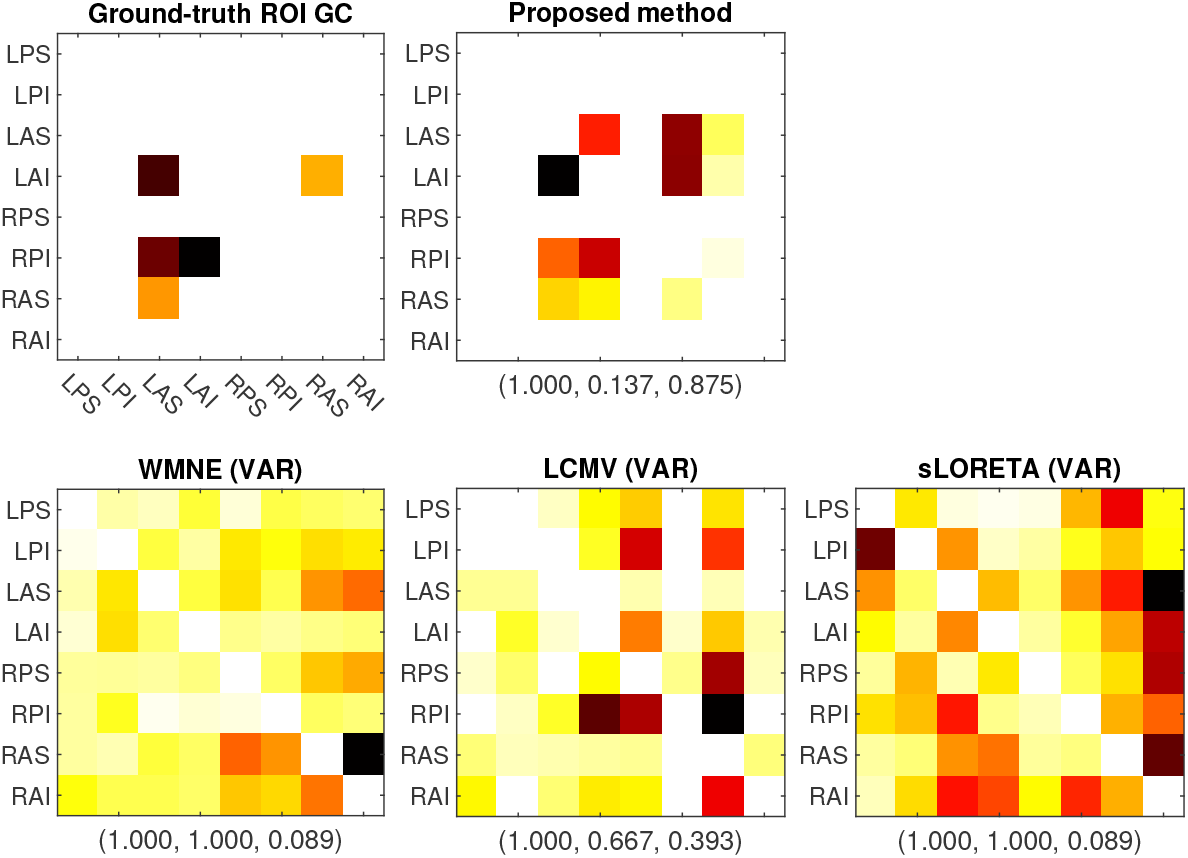
An example of estimated Granger causality. The three numbers in the parenthesis are (TPR,FPR,ACC) of the 25th instance.

**Table 1:** Averaged performance (and standard deviation) of classifying Granger causality (null versus causal).

| Indices | Proposed method | LCMV | WMNE | sLORETA |
| --- | --- | --- | --- | --- |
| TPR | 0.965 (0.12) | 0.885 (0.15) | 0.992 (0.05) | **0.997 (0.02)** |
| FPR | **0.185 (0.14)** | 0.838 (0.07) | 0.979 (0.04) | 0.994 (0.01) |
| ACC | **0.829 (0.13)** | 0.231 (0.07) | 0.114 (0.03) | 0.101 (0.02) |

## 7 Application to real EEG data

In this section, we performed an experiment on real EEG data sets and compare the findings with the previous studies that also explored brain connectivity on this data set with other methods, since the true connectivity is unknown.

### Data description

We considered a task-EEG data set containing a steady state visual evoked potential (SSVEP) EEG signals. The data were recorded from a healthy volunteer with flickering visual stimulation at 4 Hz using extended 10-20 system with 30 EEG channels. The data contained three blocks of stimulation and each of the stimulation blocks lasted 44.7 seconds. As a result, we obtained 3 trials of task-EEG segments; each of which has 11, 126 time points. The data were collected by Istanbul University, Hulusi Behcet Life Sciences Research Laboratory, Neuroimaging Unit with the approval of the local ethics committee of Istanbul University and the support of the Turkish Scientific and Technological Research Council (TUBITAK) project #108S101.

### Experiment setting

The selection of brain sources followed the details in [PLGMBB^+^18] which included the most actively ranked generators of Occipital lobe, Temporal lobe and Frontal lobe. We sample 18 sources from the six ROIs including

▪ left Occipital lobe (OL-L), right Occipital lobe (OL-R),
▪ left Temporal lobe (TL-L), right Temporal lobe (TL-R),
▪ left Frontal lobe (FL-L), right Frontal lobe (FL-R),

State-space models and source selection process were performed in node-based resolution 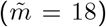.Granger causality was estimated in node-based resolution directly from model parameters. For ROI-based GC, we clustered 18 sources into 3 sources per ROI and redefined *x*_*i*_ in (26) as ROI to compute the estimated GC matrix.

Figure 10 shows active regions as the regularization parameter varies. Strongly active sources appeared in TL-R and FL-R persistently. In Figure 12, we found the second highest connection that flows from FL-L to OL-L and TL-R, and from OL-L to FL-L, OL-R, and TL-R. The linkages of exchanging information from visual cortex in occipital area to frontal lobe is known in SSVEP processing [LTZ^+^15] where they analyzed the EEG recordings using the proposed double model, compared with the partial directed coherence analysis (PDC), and also concluded this findings with previous studies using fMRI, or MEG. The three-trial estimated GC in Figure 11 shows consistently that the dominant pathways are from TL-R to regions including OL-L, TL-L, FL-L and OL-R. This is supported by the spatio-temporal analysis in [PLVHRL^+^17, PLGMBB^+^18] that the activations in frontal lobe can be preceded by stronger activations in temporal lobe but this was not found in [LTZ^+^15]. The methods in [PLGMBB^+^18] was the Hidden Gaussian Graphical State-Model (HIGGS) that relied on a frequency-domain linear state-space model with a sparse connectivity prior. The additional finding that TL played a mediator role in communication between OL and FL, reported in [PLGMBB^+^18] also agreed with previous studies in the references therein. Our framework can provide a model-based approach to confirm these active regions of exchanging information in visual processing with previous studies.

**Figure 10:**
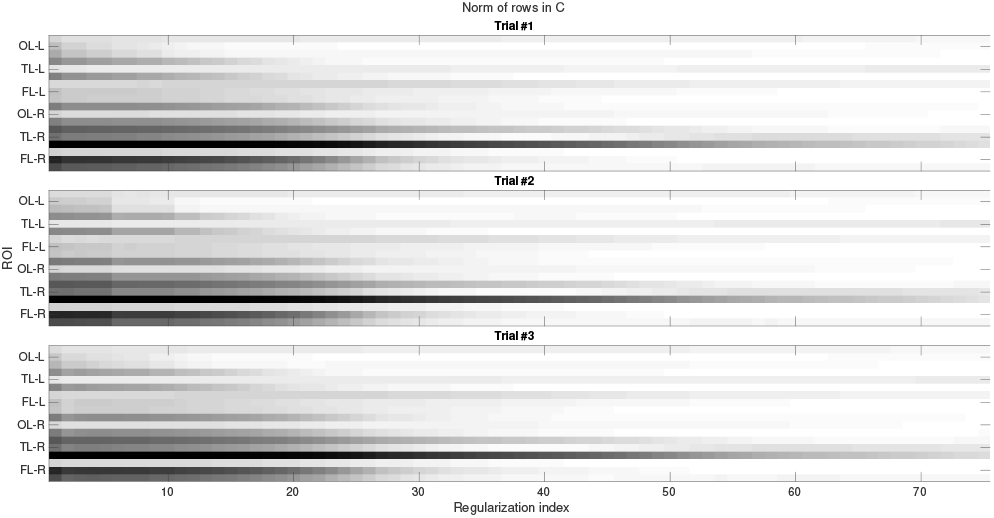
Active source selection results from SSVEP EEG data.

**Figure 11:**
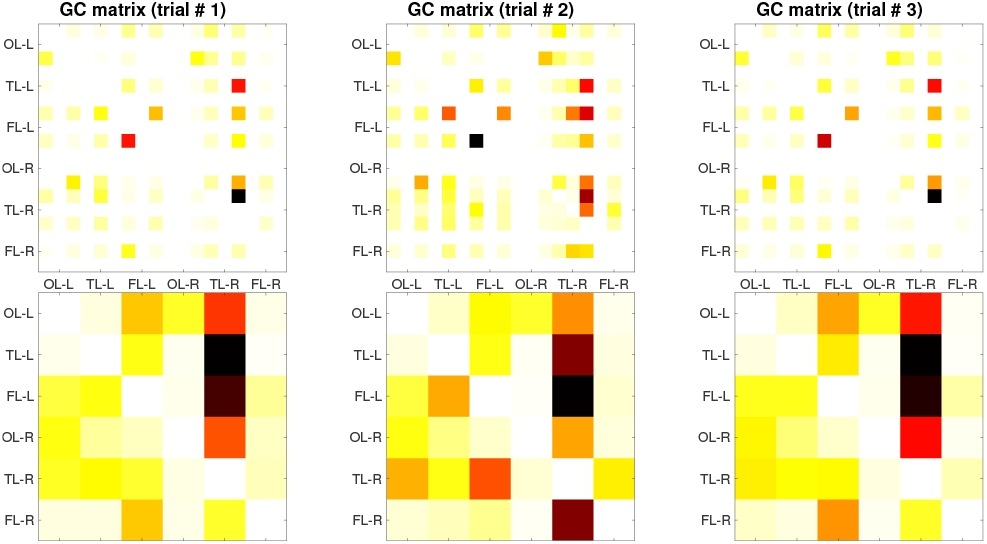
Estimated GC from three trials of SSVEP EEG data. (Top row) Node-based GC. (Bottom row) ROI-based GC.

**Figure 12:**
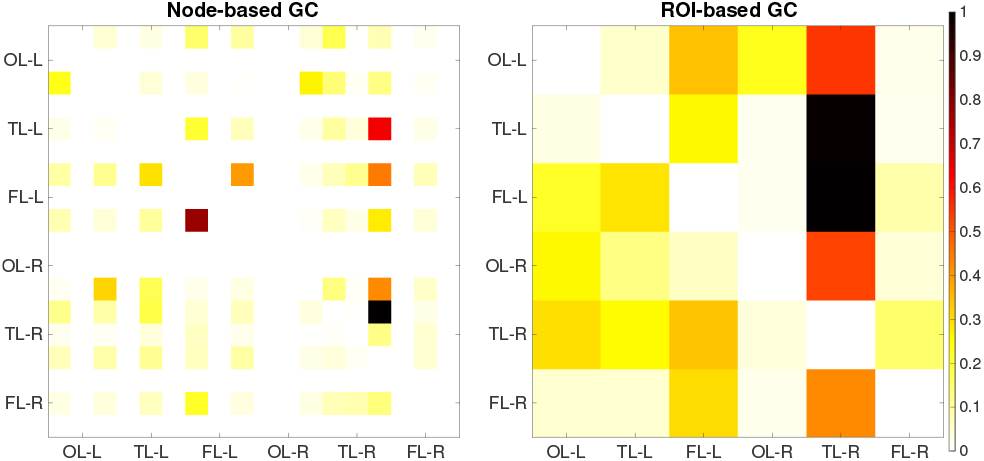
Average of estimated GC over three trials.

## 8 Limitation and concerns

Some practical concerns and limitations of the proposed method must be concluded. Firstly, the proposed method involves an approximation of *L* which needs information about sensor position, source position, and a head model. In our opinion, the latter appears to be the most uncertain parameter as subject heads can vary widely, and correspond to different head models but this information is not always measured or provided. Secondly, the overall computational complexity including model selection is high in comparison to other non-parametric approaches as the problem (31) is solved with several *γ* before using BIC to select the best model. The performance of correctly estimating the sparsity pattern in GC also relies on the choice of this model selection score, which has different finite-sample behaviors; or depends on the adopted-method of choosing the optimal *γ*, of which there are several options such as cross-validation, stability selection, or Kappa selection. Thirdly, the framework relies on a linear time-invariant state-space model, which may not entirely capture nonlinearity or non-stationarity in EEG applications with task data. Additional results shown in the supplementary material suggested that if some connections of the ground-truth network is time-varying, then our method is prone to mistakenly estimate GC as weaker than the actual strength. Nevertheless, the method can still estimate the null causality entries and time-invariant entries correctly. Also, our linear framework may fail to correctly recover GC networks when the true model is highly nonlinear, as observed in the results of false recovery and dropped ROC curves in [SGB19]. Nevertheless, from our real EEG experiment, we believe that the method yielded a certain degree of performance when the linear approximation was sufficient to capture nonlinear EEG dynamics at an operating point. Lastly, the model’s performances were evaluated and reported based on thresholding small entries of estimated GC matrices since nonparametric statistical tests such as a permutation test requires shuffling segments of the estimated source signals, which is not directly available from our approach. We believe that this step can be improved in future work when a statistical test on significant GC is well established, or by considering an alternative machine-learning framework.

## 9 Conclusion

This paper considered an estimation of linear dynamical models for EEG time series and used the model parameters to infer a causality among source signals. The model equations explain coupled dynamics of source signals and scalp EEG signals where only EEG can be measured. The definition of relationships among variables followed the idea of Granger causality (GC) that has been well-established and previously often applied on vector autoregressive (VAR) models. This work extended the VAR models to a more general class, a state-space equation, which led to a highly nonlinear characterization of GC but can be evaluated numerically via solving a discrete Riccati algebraic equation. We have provided analytical results of GC invariant properties under variations system parameters.

In order to estimate such GC matrices, we have proposed a statistical learning scheme consisting of i) subspace identification that promotes a sparse output matrix of source signal, ii) estimation of noise covariances, and iii) the estimation of GC pattern for the obtained state-space model. The subspace identification was extended by using a non-convex regularization to promote sparse rows of *C*, which can be further used to classify between active and inactive sources. The non-convex penalty was presented as a group norm of *𝓁*_2_ and *𝓁*_1/2_ to obtain a better performance than using a convex penalty in detecting zero rows of *C*. Given that the models contained equal percentages of deep and shallow sources, the overall best performance was obtained when the ground-truth model had a small portion of active sources. The median of accuracy (based on 100 runs) of classifying active sources ranged in 90 − 98%, which owes to the sparse formulation and the penalty parameter selection using BIC. When combining all the procedures and evaluating performances of estimating GC, the main factor to overall accuracy is the portion of active sources in the ground-truth models. If the true GC was sparse, the averaged accuracy was around 97% and not sensitive much to the number of electrodes and the location of sources. On the contrary, if the true GC was dense, the overall accuracy slightly degraded to 90 − 94% but had an improving trend when using less electrodes or the ground-truth system contained more deep sources. This contradicts our intuition but can be explained from the characteristic of our sparse learning framework that favors sparse models when the data samples were less available (less electrodes). The resulting sparse *C* led to sparse GC matrices which helped rule out more FPs, and hence resulted in higher accuracy.

The performance of our method was also evaluated on real SSVEP EEG data whose setting was to stimulate the human brain in the visual cortex area. Results were consistent with previous studies in the sense that a connection is found between occipital and frontal areas which are known to be related to a task of visual processing. Moreover, the temporal lobe was found to be a mediator in the connection between occipital and frontal lobes.

Some practical concerns and limitations of the proposed method can be concluded. Firstly, it requires an approximation of *L* which needs information about sensor position, source position, and a head model. In our opinion, the latter appears to be the most uncertain parameter as different subjects would correspond to different head models but this information is unlikely to be exactly known. Secondly, the overall computational complexity including model selection is high in comparison to other non-parametric approaches as the problem (31) is solved with several *γ* before using BIC to select the best model. Lastly, the reported performances were based on thresholding small entries of estimated GC matrices since nonparametric statistical test such as a permutation test requires shuffling segments of the estimated source signals, which is not directly available from our approach. We believe that this step can be improved in future work when a statistical test on significant GC is well established, or by considering a framework in machine learning.

## Acknowledgment

This research was financially supported by the Research Assistant Scholarship from Graduate School, Chulalongkorn University, and by the Chula Engineering Research grant 2019-2020. Our grateful thanks to Tamer Demiralp, Istanbul University for SSVEP EEG data set, and to Deirel Paz-Linares and Pedro A. Valdés-Sosa for providing supplemental parameters in the SSVEP EEG experiment.

## A Notations in subspace identification

We follow the notations used in [OM12] and summarize the stochastic identification method relied on the state-space model:

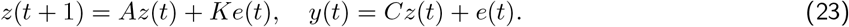

First, given output measurements 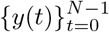, we can arrange the sequences in the following matrix.

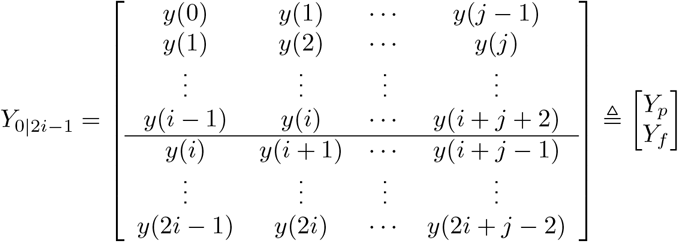

The notation *Y*_0|2*i*−1_ represents how we stack output sequences in row and columns so that it can be partitioned at the row *i* into two blocks of the past (*Y*_*p*_) and the future (*Y*_*f*_) output sequences. If *Y*_0|2*i*−1_ is row-partitioned at row *i* + 1 then it is denoted by 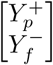. From this notation, *Y*^*i*|*I*^ is simply the output sequences starting at time *i*: *Y*_*ii*_ = *y*(*i*) *y*(*i* + 1) *y*(*i* + *j* − 1). The index *i*, which determines the number of block rows, is given by the user and should be large enough (at least larger than the maximum order of the system). The index *j* is typically chosen as *j* = *N* − 2*i* + 1 which means that all given data samples are used. In subspace identification, we often encounter the projection of row space of *A* onto the row space of *B*, denoted and given by *A/B* = *AB*^*T*^ (*BB*^*T*^)^†^*B*.

The algorithm of [OM12] involves the extended observability matrix

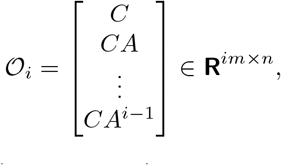

and the projection of the row space of *Y*_*f*_ (future output) on the row space of *Y*_*p*_ (past output), denoted by *ξ*_*i*_ = *Y*_*f*_ */Y*_*p*_. Given a user-defined index 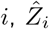 is the forward Kalman estimate of *z*(*i*) *z*(*i* + 1) *z*(*i* + *j* 1), where *j* is the time index of the last sample to be used. It was proved in Theorem 8 of [OM12] that the Kalman state sequence, 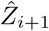, is related to the projection and the extended observability matrix via

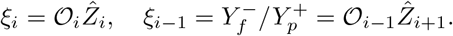

Moreover, by its definition, 𝒪 _*i* − 1_ can be obtained by stripping all block rows of 𝒪 _*i*_ except the last *r* rows. From this main result, the stochastic algorithm 3 [OM12] is described as follows.

1. Calculate the projections: *ξ*_*i*_ = *Y*_*f*_/*Y*_*p*_ and 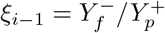.
2. Calculate the SVD of the weighted projection: 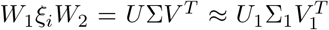 where Σ_1_ contains significantly nonzero singular values. The number of nonzero singular values determines the system order.
3. Compute 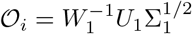 and 𝒪_*i*−1_ is obtained by extracting all rows of 𝒪_*i*_ except the last *m* rows.
4. Determine the estimated state sequences from

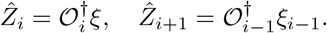

Once sequences 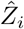 and 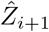 are estimated by projections, the subspace method consider the system

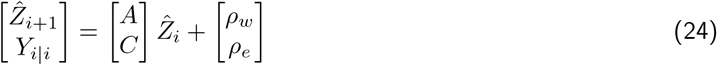

and estimate *A, C* using the least-square approach.

## B Proof of GC causality properties

This section provides a proof of Theorem 1 where we refer to the model equations:

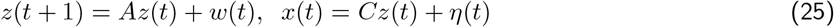

and computing a GC matrix from the definition:

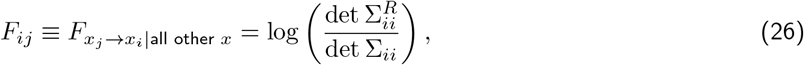

requires to solve DARE:

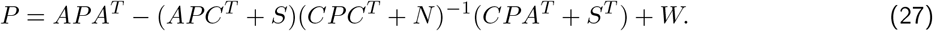

*1. A Granger causality matrix is invariant under a similarity transform*. **Proof**. A system realization of dynamical equations (25) is parameterized by (*A, C*) and the noise covariance matrices (*W, N*). Under a similarity transform to a new coordinate: 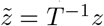, the dynamic of *x* is explained by

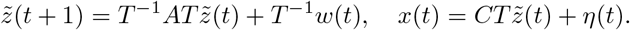

Hence, in the new coordinate of state variable, the system has another realization 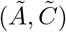 with a relation *Ã* = *T* ^−1^*AT* and 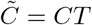. Moreover, if we define 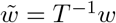, then noise covariances are

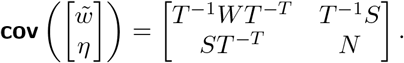

Let 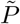 be the covariance of state estimation error in the new coordinate. It is a straightforward calculation to show that 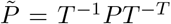 is the solution to DARE (27) and Σ is unchanged under such transformation. Hence, the Granger measure given in (26) is unchanged when it is computed using system matrices corresponding to a new coordinate system.

*2. If* 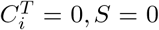 *and N is diagonal, then F*_*ij*_ = 0 *and F*_*ji*_ = 0. *As a result, the zeros of F is unchanged when N is changed under a scaling transformation*. **Proof**. We will show from the characterization of *F*_*ij*_:

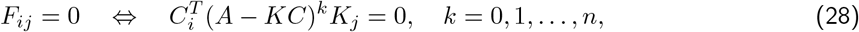

where the Kalman gain is given by *K* = (*APC*^*T*^)(*CPC*^*T*^ + *N*)^−1^ with *S* = 0. Let *i* be the index such that 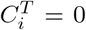. It is then obvious that the *i*th column of *APC*^*T*^ is entirely zero, *e*.*g*., (*APC*^*T*^)_*si*_ = 0 for *s* = 1, 2, *…, n*. Since *N* is diagonal, the *i*th row and the *i*th column of *CPC*^*T*^ + *N* is zero, except the (*i, i*) entry, *e*.*g*., (*CPC*^*T*^ + *N*)_*ki*_ = 0 for *k* ≠ *i* and (*CPC*^*T*^ + *N*)_*ik*_ = 0 for *k ≠i*. To see zero pattern of *CPC*^*T*^ + *N* explicitly, we have

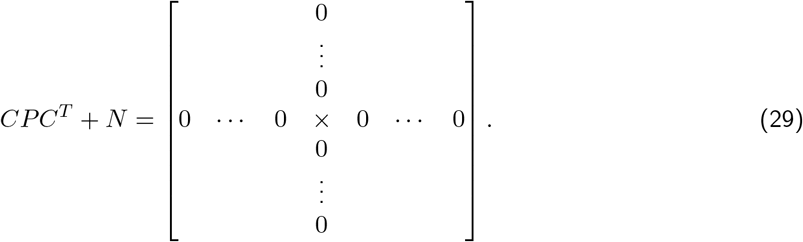

For any invertible *X*, the (*k, i*) entry of *X*^−1^ is related to the *M*_ik_ (Minor) of *X* (up to a scaling from det *X* and (−1)^*i*+*k*^). From the structure given in (29), if we remove either the *i*th row or the *i*th column, we see that *M*_*ij*_ and *M*_*ji*_ of *CPC*^*T*^ + *N* are all zero. These further imply that 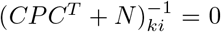for *k* ≠ *i* and 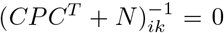 for *k* ≠ *i, e*.*g*., the *i*th row and the *i*th column of (*CPC*^*T*^ + *N*)^−1^ are entirely zero except the (*i, i*) entry. From the expression of *K*, we can conclude about its *i*th column as

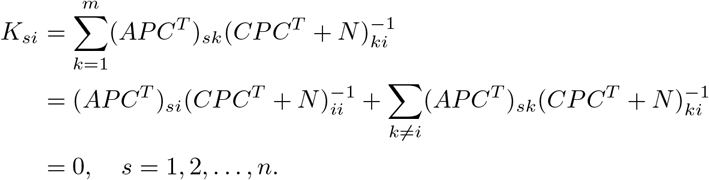

As a result, from an equivalent condition of zero GC in (28), we conclude that *F*_*ij*_ = 0 since 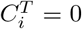 and *F*_*ji*_ = 0 because *K*_*i*_ = 0.

*3. If we permute rows of x to* 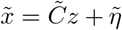, *then C is row permuted, i*.*e*., 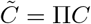 *and* 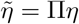 *Let* 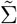 *be the covariance of* 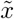. *We have* 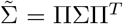. *Moreover, the GC matrix of* 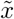 *under such permutation, is related to F by* 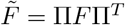. **Proof**. If we permute *x* then the noise covariances become

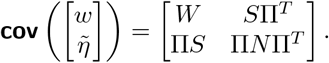

When solving DARE (27) with 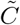 and the new covariances: *S*Π^*T*^ and Π*N* Π^*T*^, we can see that the solution *P* to DARE is unchanged. As a result, 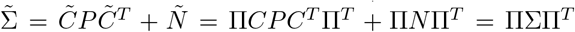, Let 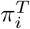 be the *i*th row of Π. Without loss of generality, let us assume that we permute *x*_*i*_ and *x*_*j*_ to be 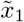 and 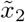 respectively. When examining a Granger cause from 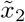 to 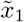, we see that 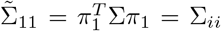and 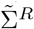 is obtained by solving DARE with the second row of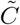 removed (equivalently, with the *j*th row of *C* removed.) Moreover, 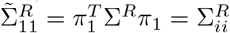. Hence, 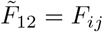. In other words, we can just permute rows and columns of *F* to obtain 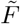.

4. *If N* = 0 *and S* = 0, *then the zero pattern of F is invariant under a scaling transformation of C*. **Proof**. It is a straightforward result when solving DARE with 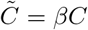that the solution *P* is unchanged if *N* = 0 and *S* = 0. Moreover, 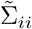 and 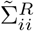 contain the same factor *β*^2^ if *N* = 0. This makes no change in the calculation of *F*_*ij*_ in (26).

## c Group-sparsity estimation of *C*

The problem of estimating *C* and *A* in

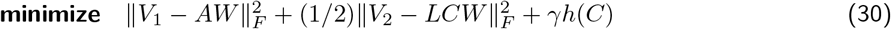

has some details of algorithm implementation. Firstly, the least-squares solution of *A* is obtained by *A* = (*V*_1_*W* ^*T*^)(*WW* ^*T*^)^−1^ and implemented by QR factorization. Secondly, for promoting a group sparsity in *C*, we use the group-norm penalty: 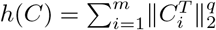 and the problem of estimating *C* is

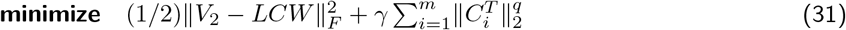

The minimization in *C*, when rearranged into a vector form, falls into a regularized least-square formulation with a group norm penalty shown as

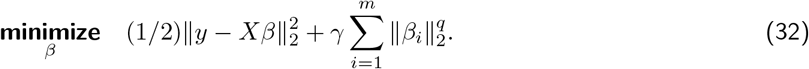

where *β* = (*β*_1_, *β*_2_, *…, β*_*m*_), *β*_*k*_ ∈ **R**^*n*^.

**Convex case**. When solving (32) with *q* = 1, it is the conventional group lasso problem, which can be solved by ADMM [PB14], or the adaptive ADMM [XFG17] that adjusts the algorithm parameter adaptively; the latter was used in our implementation. Our main formulation requires solving (31) with a series of *γ*, so an explicit bound of *γ* should be specified in order to obtain solutions *C* with all possible sparsity patterns. To this end, the bound of *γ* can be derived so that if *γ* ≥ *γ*_*c*_ then the optimal solution *C* is entirely zero. We derive the zero-subgradient condition for (31) when *q* = 1.

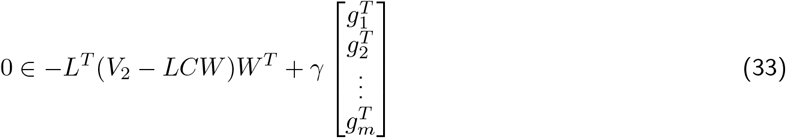

where 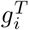 is a subgradient of 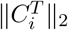 with a known property that 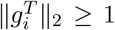 for all *i* if *C* = 0. If *C* = 0 at optimum, then (33) reduces to

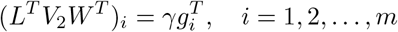

where (*L*^*T*^ *V*_2_*W* ^*T*^)_*i*_ is the *i*th row of *L*^*T*^ *V*_2_*W* ^*T*^. Since 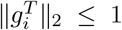 for all *i*, when *C* = 0, it follows that *γ* ≥ max_*i*=1,2,…,m_ ∥(*L*^*T*^ *V*_2_*W* ^*T*^)_*i*_∥_2_. The critical value *γ*_*c*_ for (31) is therefore

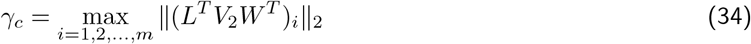

which depends only the problem parameters *L, V*_2_, *W* and can be computed beforehand.

**Nonconvex case**. When solving (31) with *q* = 1*/*2, we split the objective function to *F* (*C*) = *f* (*C*) + *γh*(*C*) where 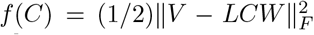 and 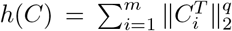, and applied the nmAPG algorithm proposed in [LL15]. This requires the proximal operator of *q*-norm and the Lipschitz constant of ∇_*c*_*f* which is *L*^*T*^ (*V LCW*)*W* ^*T*^. The algorithm can be described as follows.

### Algorithm 1

nmAPG algorithm [LL15] for solving (31)

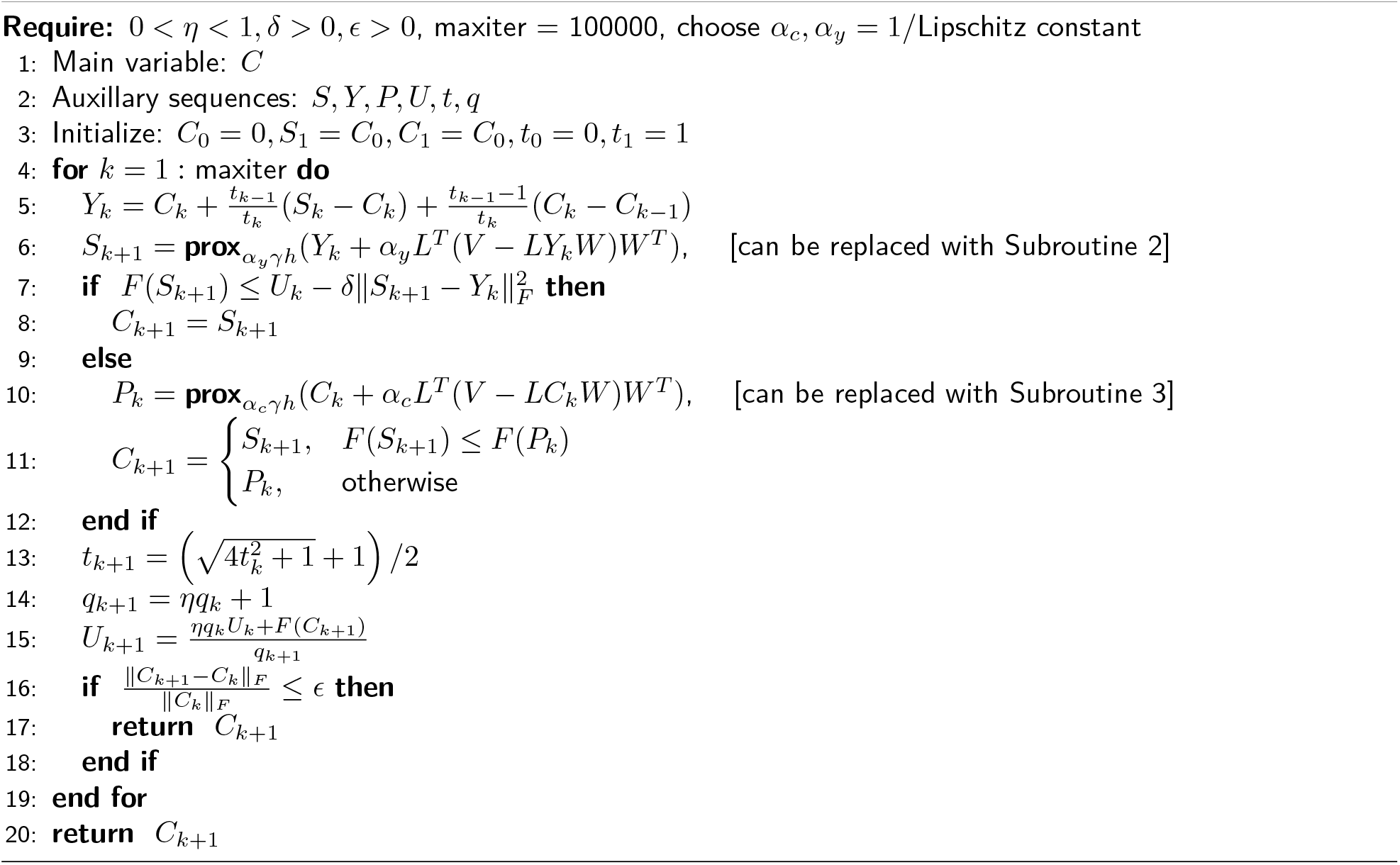

We set parameters: *δ*= 0.7, *η* = 0.9, *ϵ* = 10^−5^ in our implementation. An initialized solution *C*_0_ in line 3 can be other choices such as the least-square solution (if valid). When solving the problem for several values of *γ*, we can initialize *C*_0_ as the optimal *C* returned from solving the problem with the previous value of *γ*. The required constant of the algorithm is the Lipschitz constant of ∇_*c*_*f*. This is then obtained from the following inequality:

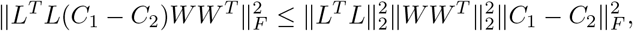

where we denote ∥ · ∥_2_ as the spectral norm and ∥ · ∥_*f*_ as the Frobenius norm of a matrix. We also have used the fact that 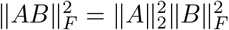. As a result, the Lipschitz of ∇_*c*_*f* is given by 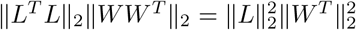, and used in a step size selection in the proximal step.

As for the proximal operator of *h*, recall its definition as **prox** (*V*) = argmin_*C*_ 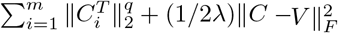. Denote 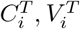 for *i* = 1, *…, m*, the *i*th row of *C* and *V*, respectively. Since *h*(*C*) is separable in each row of *C*, **prox**_*λh*_(*V*) consists of *m* rows; each of which can be computed in parallel as

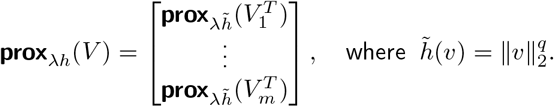

For *q* = 1*/*2, the function 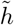 has the closed-form proximal operator given by [HLM^+^17].

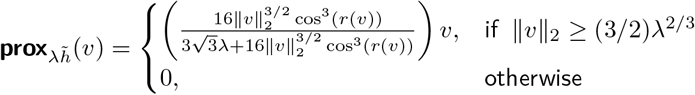

with *r*(*v*) = *π/*3 − arccos 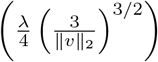When the Lipschitz constant of ∇*f* is really large, the step sizes *α*_*c*_, *α*_*y*_ can be very small. Then, the proximal steps in line 6 and 10 can be optionally replaced by the Barzillai-Borwein (BB) line search to acheive a possible larger step. The following routines are referred from Supplementary Material of [LL15].

### Subroutine 2

Barzillai-Borwein line search for *α*_*y*_

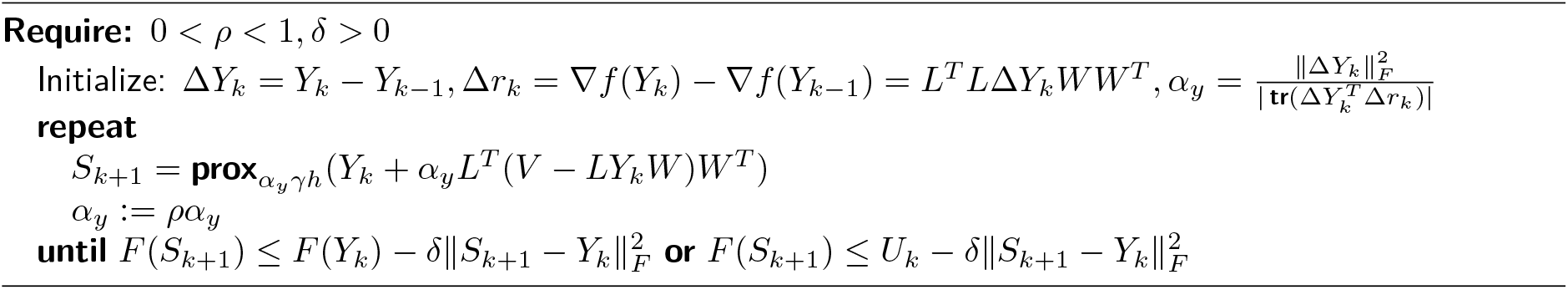

### Subroutine 3

Barzillai-Borwein line search for *α*_*c*_

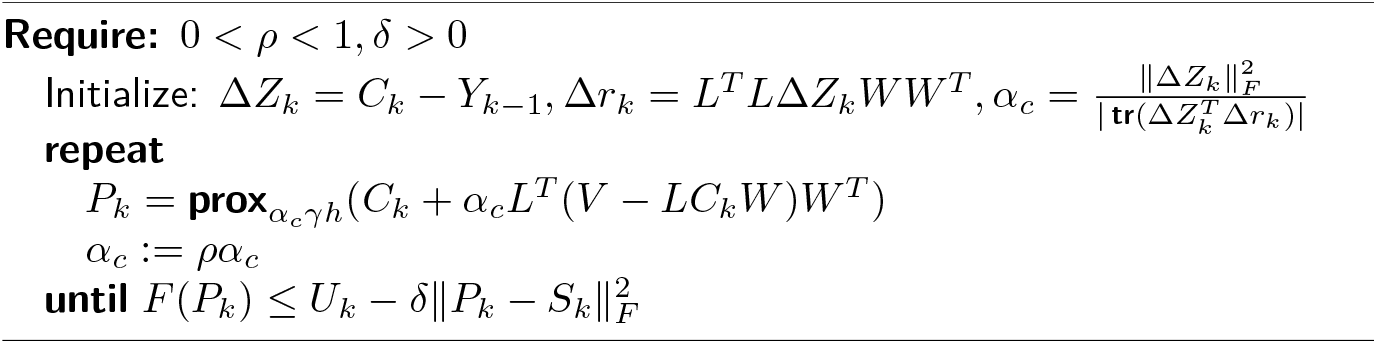

Note that, in the implementation of solving (31), we perform QR factorizations on problem parameters as *V*_2_ = *R*_*v*_*Q*^*T*^ and *W* = *R*_*w*_*Q*^*T*^. Therefore, the problem (31) is equivalent to

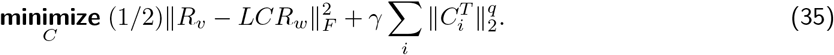

To find a range of *γ* for using with (31) to extract different sparsity levels of *C*, we use a property that the non-convex penalty (when *q <* 1) often gives sparser solutions than the convex penalty (when *q* = 1) given the same *γ* [HTW15]. *Hence, we use the same γ*_*c*_ given in (34) when solving the problem with *q* = 1*/*2.

## *D 𝓁*_2_-regularized estimation of *C*

In contrast to sparseness property of the solutions when using *𝓁_p,q_* penalty, the estimation of *C* in (30) with the *𝓁*_2_-regularization: 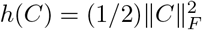 is the problem:

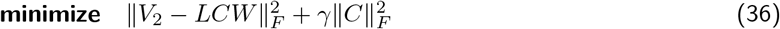

which is known to gives a typically dense solution. However, as a norm penalty, using *h*(*C*) also forces the solution to zero when *γ* is large. We will show explicitly that as *γ* → ∞, more weight is penalized on the norm of *C* and hence, *C* → 0. The optimality condition is the Sylvester equation in *C*:

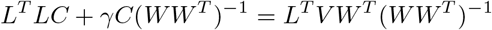

that can be arranged in the form: *AC* + *CB*(*γ*) = *F* where *A* = *L*^*T*^ *L, B*(*γ*) = *γ*(*WW* ^*T*^)^−1^, and *F* = *L*^*T*^ *V W* ^*T*^ (*WW* ^*T*^)^−1^. This equation can be vectorized to a form of *M* (*γ*)*z* = *b* where *z* = **vec**(*C*) and has a unique solution: *z* = *M* ^−1^(*γ*)*b*, since *A* and −*B*(*γ*) have no common eigenvalues. We will show that *z* → 0 as *γ* → ∞. It can be shown that *M* can be represented as

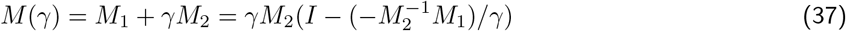

where *M*_1_ = *I*_*n*_ ⊗ *A* and *M*_2_ = *B*^*T*^ ⊗ *I*_*m*_. If we define 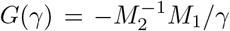 and if 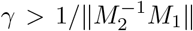 then∥*G*(*γ*) ∥ _2_ *<* 1 and (*I*− *G*(*γ*))^−1^ can be expanded by the geometric series: 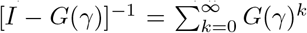 As a result, by properties of norm,

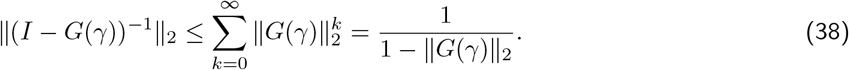

From (37) and (38), it follows that

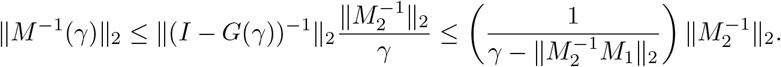

Hence, if *γ* → ∞ then ∥*M* ^−1^(*γ*) ∥_2_ → 0 and that ∥*z* ∥≤ ∥*M* ^−1^(*γ*) ∥ _2_ ∥*b* ∥≤0. The solution *z* = **vec** *C* converges to zero as the penalty parameter *γ* approaches infinity.

## E Noise covariance estimation

Consider the reduced noise estimation problem:

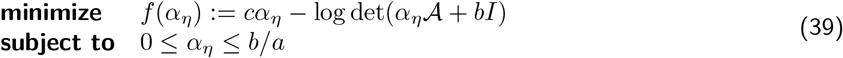

where the optimization variable *α*_*η*_ is the uniform variance of noise *η*. Refer to the original covariance estimation problem presented in the paper. Once, *α*_*η*_ is determined, we can solve the uniform variance of noise *v* from *α*_*v*_ = −*aα*_*η*_ + *b*. We show in this section that the solution *α*_*η*_ can be obtained in almost closed-form. The zero-gradient condition of the objective function is

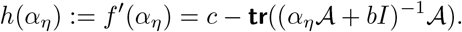

If the critical point of *f*, denoted as 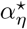 in Figure 13, is already in the interval (0, *b/a*) then the solution of (39) is just obtained by solving *f* ^′^(*α*_*η*_) = 0 (by any numerical methods such as a bisection). If 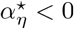, then minimizing *f* over the interval [0, *b/a*] returns 0 as the optimal solution. In the last case, if 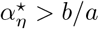 then *f* has the minimum value on the interval [0, *b/a*] at *b/a*. Therefore, we conclude that the solution of the problem (39) can be obtained from one of the following three cases.

**Figure 13:**
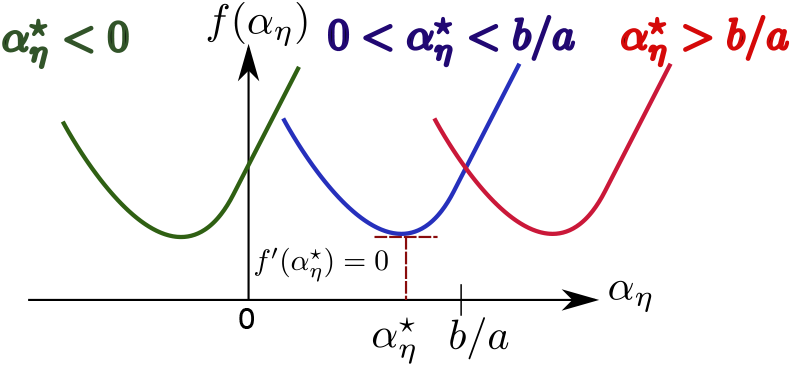
The problem of estimating *α*_*η*_ in (39) has three cases.

1. If *f* ^′^(0) *<* 0 and *f* ^′^(*b/a*) *>* 0 then find the zero of *h*(*α*_*η*_) using the bisection method.
2. If *f* ^′^(0) *>* 0 and *f* ^′^(*b/a*) *>* 0 then *α*_*η*_ = 0 and *α*_*v*_ = **tr**(Σ_*e*_)*/r*.
3. If *f* ^′^(0) *<* 0 and *f* ^′^(*b/a*) *<* 0 then *α*_*η*_ = *b/a* = **tr**(Σ_*e*_)*/* **tr**(*LL*^*T*^) and *α*_*v*_ = 0.

## F Time-varying ground-truth GC

This additional experiment is aimed to illustrate the method performance when the framework assumptions are not held. The proposed source-EEG model is a set of linear time-invariant state-space equations. Recall the data generation method explained in Section V of the main paper. Ground-truth GC structures, generated from our scheme, have the same pattern as that of the VAR coefficients of VARMA process explaining the source dynamics. Therefore, we would like to generalize the ground-truth GC pattern to be time-varying due to the changes in VAR coefficients. While it is likely that the overall performance can be degraded, it is more interesting to explore which sub-procedure that mainly affects the performance. For this purpose, we firstly consider a state-space GC estimation without involving any EEG dynamics to see how the time-invariant GC estimation is sensitive to the time-varying ground-truth structure. Secondly, when the EEG observation equation is combined with the state-space source dynamics, we investigate how the process of estimating *C* in the source equation affects the GC estimation performance.

To answer the first question, we generated a VARMA process with the VAR coefficient of the form 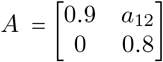 and the diagonal filter that introduces the MA part is given by *G*(*z*) = **diag**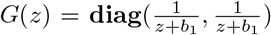. Hence the system *H*(*z*) = *G*(*z*)*A*(*z*)^−1^ = *G*(*z*)(*I* — *Az*^−1^) represents the state-space model: *z*(*t* + 1) = A *z*(*t*) + *Bw*(*t*), *x*(*t*) = *Cz*(*t*). From the structure of *A*, and the diagonal property of *G*, we can read that *x*_1_ does not have a Granger cause to *x*_2_ and *x*_2_ has a Granger cause to *x*_1_ as long as *a*_12_ ≠ 0. The value of *a*_12_ signifies the ground-truth GC pattern, so we vary *a*_12_ for *J* times and obtain *J* +1 switching sub-systems: *H*_1_(*z*), *…, H*_J+1_(*z*). At *t* = 0, the system is governed by *H*_1_, and then switched to *H*_2_ at time *t*_1_, and so on until the last transition time. The generated time series of *x*(*t*) with time points of *T* = 1000 can then be divided into *J* stationary segments. However, the GC estimation of *x*(*t*) is performed on the whole non-stationary data, violating the method assumption. We generate 500 runs of *x*(*t*) and obtain 500 samples of GC estimates, 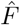.

The result in Figure 14 is obtained when *a*_12_ decreases from 0.8 to 0 with *J* = 1, 4, 19. This corresponds to the ground-truth GC matrix, *F* varying from 0.7 to 0 (approximately). The 500-samples of 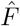 have the histogram concentrated around the interval (0.1, 0.3). It means that the method would misclassify that *x*_2_ has a Granger cause to *x*_1_. These unfortunate results do not appear to be improved even when *a*_12_ is more slowly varying (*J* = 19). From a brain connectivity application point of view, if the framework is performed on task data containing a mix of resting and stimulated states, where the true connectivity switches from active to inactive, then the estimated connectivity is fused toward causal.

**Figure 14:**
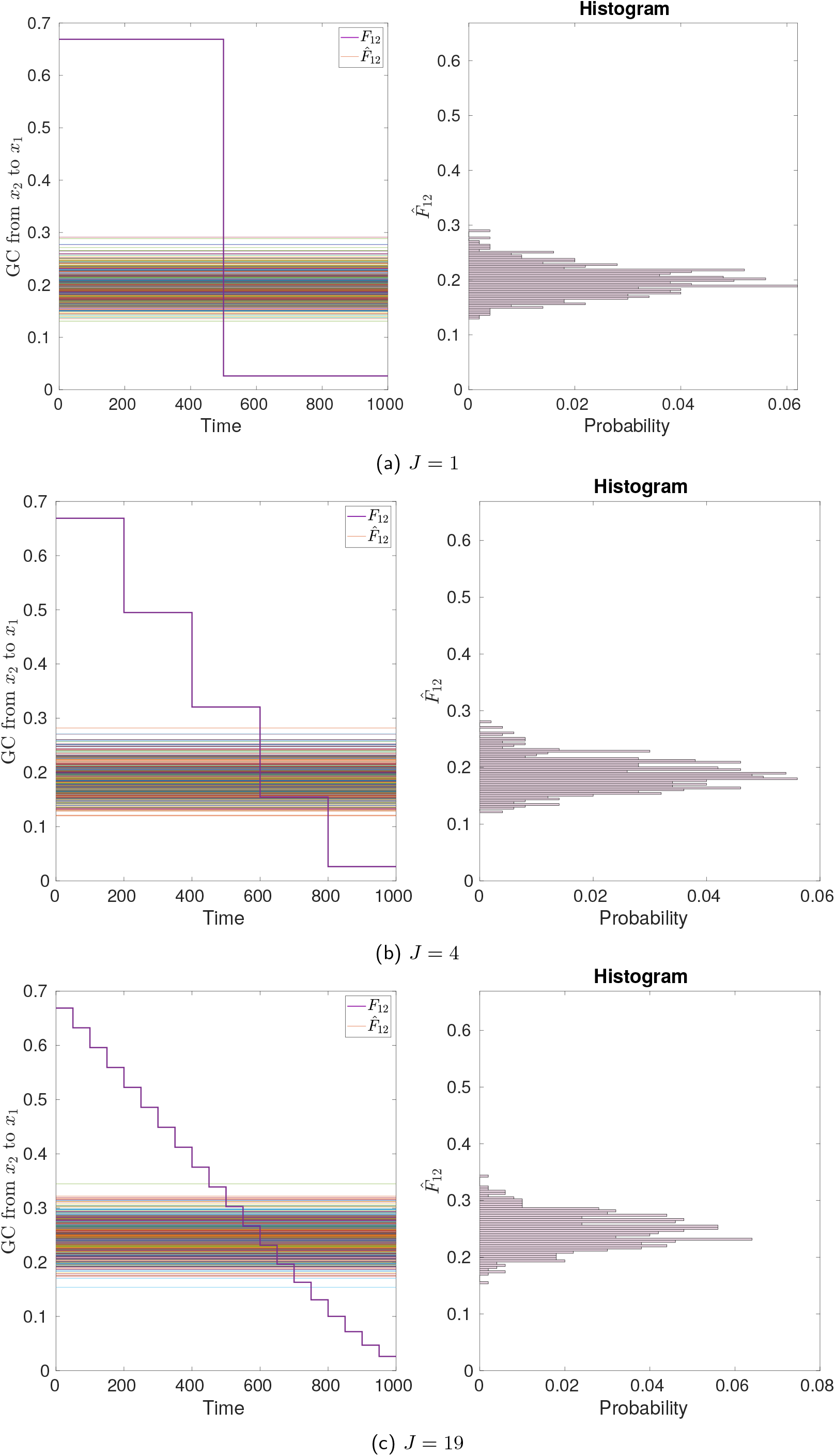
Time-varying ground-truth GC of **state-space equation** and its time-invariant estimates from 500 runs. The ground-truth GC varies in a **wide** range.

However, when *a*_12_ varies from 0.8 to 0.6 (smaller range), Figure 15 shows that 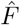 stays in the causal range (0.4, 0.6) but this is biased toward a lower value than the actual one in *F*. As a brain connectivity interpretation, if the brain network from one region to another has become weaker (but not completely gone), then our method still has a chance of detecting such linkage, while the estimated strength does not correctly represent the ground-truth magnitude.

**Figure 15:**
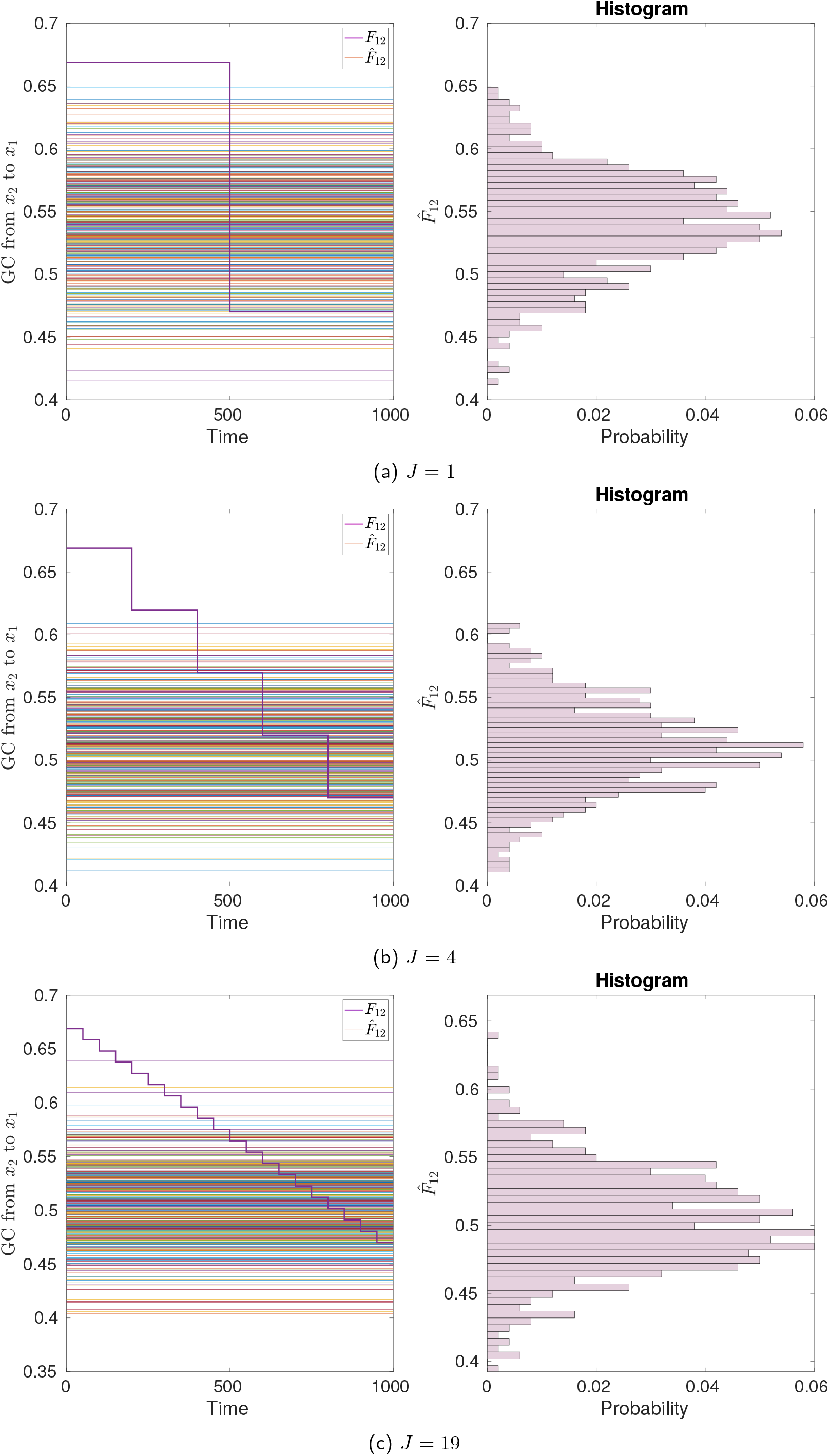
Time-varying ground-truth GC of **state-space equation** and its time-invariant estimates from 500 runs. The ground-truth GC varies in a **small** range.

As seen, even when estimating GC of a state-space model alone can lead to a bias, so our answer to the second question might not be positive. Here, we have added the EEG observation equation: *y*(*t*) = *Lx*(*t*) + *e*(*t*) where *L* is known. We have used 3-dimensional VAR part of lag 2. In the ground-truth system, there are 2 inactive and 3 active sources, and the active ones have GC structure that *F*_21_ is fixed positive, *F*_12_ = *F*_31_ = *F*_32_ = 0 while *F*_13_ and *F*_23_ are allowed to be varied *J* times. From 500 runs of *y*(*t*);each of which has 5000 time points, we apply our framework to estimate state-space parameters, classify inactive sources, and estimate *C*. Since 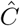 estimated from a regularized method is known to be bias, we also hypothesize that the overall performance of GC estimation is less accurate.

Each subfigure in Figure 16 has left and right *y*-axes. The blue histogram of 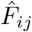 is read with the left *y*-axis while the vertical red line shows how the ground-truth *F*_*ij*_ varies with time, read with the right *y*-axis. The estimation result of the null GC entries (*F*_12_, *F*_31_, *F*_32_) and the time-invariant causal entries (*F*_21_) are robust to time-varying changes of other entries in the ground-truth. Considering the causal time-varying GC entries (*F*_13_, *F*_23_), we see the histograms shifted toward the lower range of (0.01, 0.04) when the actual GC decreases from 0.1 to 0 (approximately), regardless of the GC change rate (*J* = 1, 9). It shows that our method may misclassify those time-varying GC entries as null with some significant probability. We explore further if the direction of varying GC has some influence on the result, we found that the method still introduces a bias toward a lower range even if the true GC is increased from 0 to 0.1 (graphs are not shown). To conclude from this experiment, when some connections of the underlying causality structure have changed, which violate our framework assumption, our method is prone to mistakenly estimate GC as weaker than the actual strength. On the plus side, the method can still estimate the zero entries and time-invariant entries correctly.

**Figure 16:**
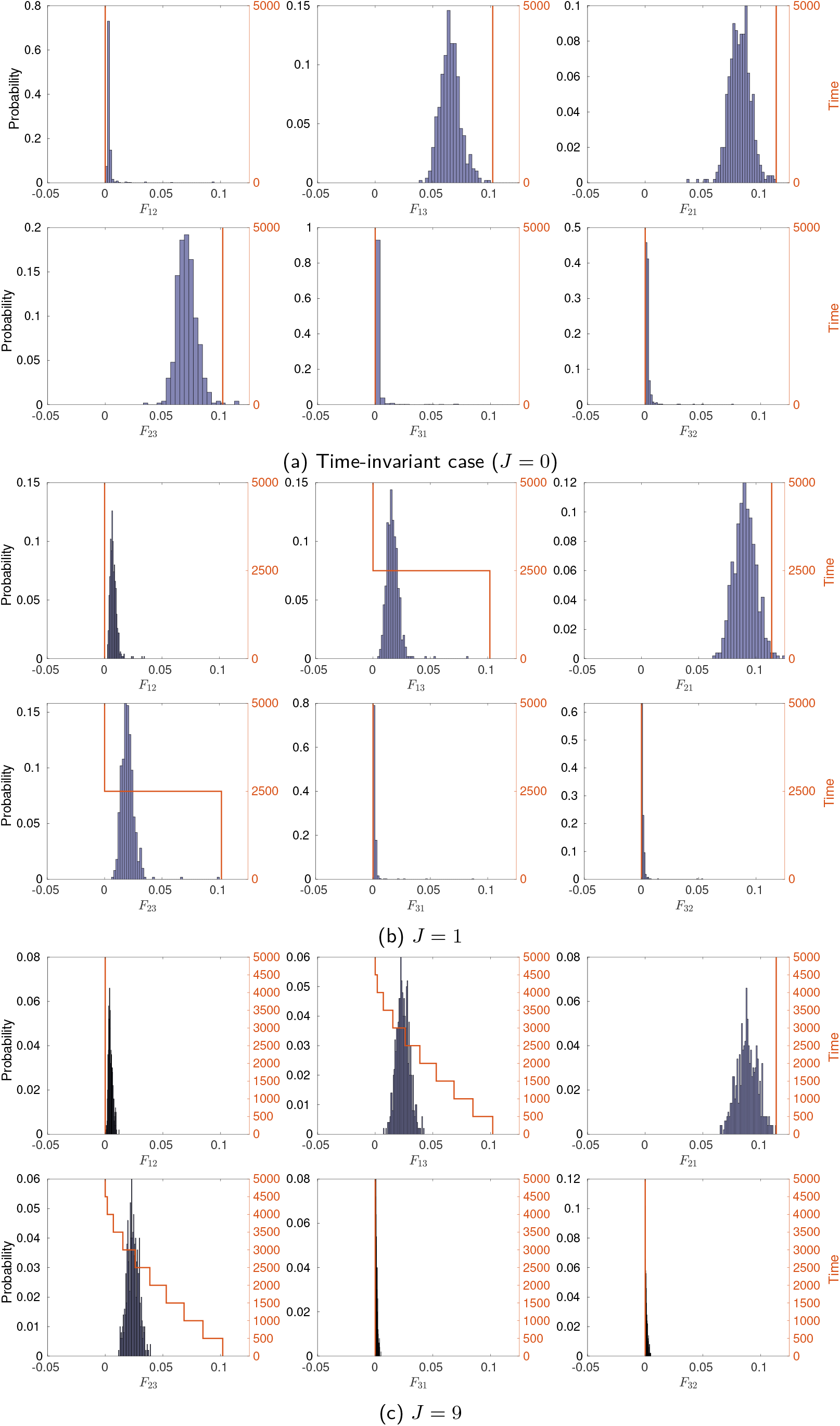
Time-varying ground-truth GC of **source-EEG equations** and its time-invariant estimates from 500 runs.

## References

[ACM+07] L. Astolfi, F. Cincotti, D. Mattia, M. G. Marciani, L. A Baccala, F. de Vico Fallani, S. Salinari, M. Ursino, M. Zavaglia, L. Ding, et al. Comparison of different cortical connectivity estimators for high-resolution EEG recordings. Human brain mapping, 28(2):143–157, 2007.

[APS+19] A. Anzolin, P. Presti, F. Van De Steen, L. Astolfi, S. Haufe, and D. Marinazzo. uantifying the effect of demixing approaches on directed connectivity estimated between reconstructed EEG sources. Brain Topography, 32(4):655–674, 2019.

[BS11] L. Barnett and A.K. Seth. Behaviour of Granger causality under filtering: theoretical invariance and practical application. Journal of Neuroscience Methods, 201(2):404–419, 2011.

[BS14] L. Barnett and A. K. Seth. The MVGC multivariate Granger causality toolbox: a new approach to Granger-causal inference. Journal of Neuroscience Methods, 223:50–68, 2014.

[BS15] L. Barnett and A. K. Seth. Granger causality for state-space models. Physical Review E, 91(4):1– 6, 2015.

[CGHJ12] J. Casals, A. García-Hiernaux, and M. Jerez. From general state-space to VARMAX models. Mathematics and Computers in Simulation, 82(5):924–936, 2012.

[CRTVV10] Bing Leung Patrick Cheung, Brady Alexander Riedner, Giulio Tononi, and Barry D Van Veen. Estimation of cortical connectivity from EEG using state-space models. IEEE Transactions on Biomedical Engineering, 57(9):2122–2134, 2010.

[CWM12] J. Chiang, Z. Jane Wang, and M. J. McKeown. A generalized multivariate autoregressive (GMAR)-based approach for EEG source connectivity analysis. IEEE Transactions on Signal Processing, 60(1):453–465, 2012.

[dSFK+16] F. Van de Steen, L. Faes, E. Karahan, J. Songsiri, P.A. Valdes-Sosa, and D. Marinazzo. Critical comments on EEG sensor space dynamical connectivity analysis. Brain Topography, pages 1–12, 2016.

[GB19] A.J. Gutknecht and L. Barnett. Sampling distribution for single-regression Granger causality estimators. arXiv preprint arXiv:1911.09625, 2019.

[GHAEC08] G. Gómez-Herrero, M. Atienza, K. Egiazarian, and J. L. Cantero. Measuring directional coupling between EEG sources. NeuroImage, 43(3):497–508, 2008.

[GPO12] R. E. Greenblatt, M. E. Pflieger, and A. E. Ossadtchi. Connectivity measures applied to human brain electrophysiological data. Journal of Neuroscience Methods, 207(1):1–16, 2012.

[Hau12] S. Haufe. Towards EEG Source Connectivity Analysis. PhD thesis, Technische Universität Berlin, Germany, 2012.

[HAVS+19] B. He, L. Astolfi, P.A. Valdés-Sosa, D. Marinazzo, S.O. Palva, C. Bénar, C.M. Michel, and T. Koenig. Electrophysiological brain connectivity: theory and implementation. IEEE Transactions on Biomedical Engineering, 66(7):2115–2137, 2019.

[HBCN+17] L. Albera H. Becker, P. Comon, J.-C. Nunes, R. Gribonval, J. Fleureau, P. Guillotel, and I. Merlet. Sissy: An efficient and automatic algorithm for the analysis of EEG sources based on structured sparsity. NeuroImage, 157:157–172, 2017.

[HE16a] S. Haufe and A. Ewald. A simulation framework for benchmarking EEG-based brain connectivity estimation methodologies. Brain Topography, pages 1–18, 2016.

[HE16b] S. Haufe and A. Ewald. A simulation framework for benchmarking EEG-based brain connectivity estimation methodologies. Brain Topography, pages 1–18, 2016.

[HJ13] R. A. Horn and C. R. Johnson. Matrix Analysis. Cambridge university press, 2nd edition, 2013.

[HLM+17] Y. Hu, C. Li, K. Meng, J. Qin, and X. Yang. Group sparse optimization via lp,q regularization. Journal of Machine Learning Research, 18(30):1–52, 2017.

[HNMN13] S. Haufe, V. V Nikulin, K. Müller, and G. Nolte. A critical assessment of connectivity measures for EEG data: a simulation study. NeuroImage, 64:120–133, 2013.

[HPH16] Y. Huang, L.C. Parra, and S. Haufe. The New York Head: A precise standardized volume conductor model for EEG source localization and tES targeting. NeuroImage, 140:150–162, 2016.

[HTF09] T. Hastie, R. Tibshirani, and J. Friedman. The Elements of Statistical Learning: Data Mining, Inference and Prediction. Springer, 2nd edition, 2009.

[HTN+10] S. Haufe, R. Tomioka, G. Nolte, K. Müller, and M. Kawanabe. Modeling sparse connectivity between underlying brain sources for EEG/MEG. IEEE Transactions on Biomedical Engineering, 57(8):1954–1963, 2010.

[HTW15] T. Hastie, R. Tibshirani, and M. Wainwright. Statistical learning with sparsity: the lasso and generalizations. Chapman and Hall/CRC, 2015.

[LL15] H. Li and Z. Lin. Accelerated proximal gradient methods for nonconvex programming. In Advances in Neural Information Processing Systems 28, pages 379–387, 2015.

[LTZ+15] F. Li, Y. Tian, Y. Zhang, K. Qiu, C. Tian, W. Jing, T. Liu, Y. Xia, D. Guo, D. Yao, et al. The enhanced information flow from visual cortex to frontal area facilitates SSVEP response: evidence from model-driven and data-driven causality analysis. Scientific Reports, 5(1):1–11, 2015.

[Lüt05] H. Lütkepohl. New Introduction to Multiple Time Series Analysis. Springer, 2005.

[LWVS15] X. Lei, T. Wu, and P. Valdes-Sosa. Incorporating priors for EEG source imaging and connectivity analysis. Frontiers in Neuroscience, 9(284):1–12, 2015.

[MMARPH14] J. Montoya-Martínez, A. Artés-Rodríguez, M. Pontil, and L. K. Hansen. A regularized matrix factorization approach to induce structured sparse-low-rank solutions in the EEG inverse problem. EURASIP Journal on Advances in Signal Processing, 19:97, 2014.

[OM12] P. Van Overschee and B. De Moor. Subspace identification for linear systems: Theory– Implementation–Applications. Springer Science & Business Media, 2012.

[PB14] N. Parikh and S. Boyd. Proximal algorithms. Foundations and Trends in Optimization, 1(3):127– 239, 2014.

[PiS18] N. Plub-in and J. Songsiri. State-space model estimation of EEG time series for classifying active brain sources. In 2018 11th Biomedical Engineering International Conference, pages 1–5. IEEE, 2018.

[PiS19] N. Plub-in and J. Songsiri. Estimation of Granger causality of state-space models using a clustering with Gaussian mixture model. In Proceedings of IEEE International Conference on Systems, Man, and Cybernetics (IEEE SMC). IEEE, 2019.

[PLGMBB+18] D. Paz-Linares, E. Gonzalez-Moreira, J. Bosch-Bayard, A. Areces-Gonzalez, M.L. Bringas-Vega, and P.A. Valdes-Sosa. Neural connectivity in M/EEG with hidden hermitian Gaussian graphical model. arXiv:1810.01174 [stat.ME], pages 1–34, 2018.

[PLVHRL+17] D. Paz-Linares, M. Vega-Hernandez, P.A. Rojas-Lopez, P.A. Valdes-Hernandez, E. Martínez-Montes, and P.A. Valdes-Sosa. Spatio temporal EEG source imaging with the hierarchical bayesian elastic net and elitist lasso models. Frontiers in Neuroscience, 11:635, 2017.

[PS16] A. Pruttiakaravanich and J. Songsiri. A Review on Exploring Brain Networks from fMRI Data. Engineering Journal, 20(3):1–28, 2016.

[SC13] S. Sanei and J. A. Chambers. EEG Signal Processing. John Wiley & Sons, 2013.

[SGB19] Y. Shen, G.B. Giannakis, and B. Baingana. Nonlinear structural vector autoregressive models with application to directed brain networks. IEEE Transactions on Signal Processing, 67(20):5325– 5339, 2019.

[STOS17] S.B. Samdin, C.M. Ting, H. Ombao, and S.H. Salleh. A unified estimation framework for state-related changes in effective brain connectivity. IEEE Transactions on Biomedical Engineering, 64(4):844–858, 2017.

[SWF13] W. Sun, J. Wang, and Y. Fang. Consistent selection of tuning parameters via variable selection stability. The Journal of Machine Learning Research, 14(1):3419–3440, 2013.

[SYW+16] A. Sohrabpour, S. Ye, G.A. Worrell, W. Zhang, and B. He. Noninvasive electromagnetic source imaging and granger causality analysis: an electrophysiological connectome (econnectome) ap-proach. IEEE Transactions on Biomedical Engineering, 63(12):2474–2487, 2016.

[TBM+11] F. Tadel, S. Baillet, J. C. Mosher, D. Pantazis, and R. M Leahy. Brainstorm: a user-friendly application for MEG/EEG analysis. Computational Intelligence and Neuroscience, page 8, 2011.

[WTO16] Y. Wang, C. Ting, and H. Ombao. Modeling effective connectivity in high-dimensional cortical source signals. IEEE Journal of Selected Topics in Signal Processing, 10(7):1315, 2016.

[XFG17] Z. Xu, M. Figueiredo, and T. Goldstein. Adaptive ADMM with spectral penalty parameter selection. In Aarti Singh and Jerry Zhu, editors, Proceedings of the 20th International Conference on Artificial Intelligence and Statistics, volume 54 of Proceedings of Machine Learning Research, pages 718–727. PMLR, Apr 2017.

[YYR16] M. Tarr Y. Yang, E. Aminoff and K. E Robert. A state-space model of cross-region dynamic connectivity in MEG/EEG. Advances in Neural Information Processing Systems 29, pages 1234– 1242, 2016.

